# AgamOBP4-targeted drug repurposing identifies novel mosquito repellent scaffolds

**DOI:** 10.64898/2026.10.21.758783

**Authors:** Jorge-R. Alonso-Fernández, Evgenia C.V. Stamati, Christina E. Drakou, Maria Mazavli, Vasileios Karras, Dimitrios P. Papachristos, Antonios Michaelakis, Spyros E. Zographos, Horacio Pérez-Sánchez, Katerina E. Tsitsanou

## Abstract

Odorant-binding proteins (OBPs) play central roles in insect chemoreception and represent attractive molecular targets for the discovery of novel behavior-modifying compounds. We report two crystal structures of *Anopheles gambiae* OBP4: one complexed with the fluorescent probe N-phenyl-1-naphthylamine and a second containing two indole molecules simultaneously bound within the ligand-binding cavity. Structural comparison revealed two binding subsites accommodating the aromatic moieties of 1-NPN or two indole molecules, while identifying Thr58 and Thr70 as key residues in ligand recognition. A structure-based virtual screening campaign was performed against a volatility-filtered library from DrugBank, FooDB, BioNet, and natural-volatiles datasets. Twenty-three ligands were selected for fluorescence competitive binding assays, identifying four AgamOBP4 binders with micromolar affinities (*K*i = 2.4-21.8 μM). Among them were benzoyl peroxide, an active anti-acne ingredient, Avex-73 (blarcamesine), a clinical-stage drug candidate, and 2-phenylethyl benzoate, a food- and fragrance-associated ester. Additional candidates include o,p′-DDE, a metabolite of the insecticide DDT, and fluorene, a polycyclic aromatic hydrocarbon detected in foods as environmental or processing-derived contaminants. Seven selected compounds were subsequently evaluated against *Aedes albopictus* in *in-vivo* behavioral assays, with benzoyl peroxide displaying the highest repellency (RI = 81.8% at 0.2 mg cm^-2^). AI-based protein-ligand modeling provided insights into ligand recognition and suggested relationships among binding mode, binding affinity, and bioactivity. Collectively, these findings establish the structural basis of ligand recognition by AgamOBP4, highlight the potential of drug-repurposing and food-derived chemical space as sources of next-generation mosquito repellents, and open new opportunities for the development of OBP4-based biosensors for food-quality and environmental monitoring applications.

**Highlights:**

- Novel AgamOBP4 crystal complexes revealed a dual aromatic-recognition binding site
- Structure-based virtual screening of >139,000 compounds from DrugBank, FooDB, and BioNet uncovered novel AgamOBP4 ligands from drug-repurposing and food-derived chemical spaces
- *In vitro* validation identified the clinical-stage candidate Avex-73 and the anti-acne agent benzoyl peroxide as novel low-micromolar AgamOBP4 ligands.
- Binding of the DDT metabolite o,p′-DDE suggests a possible involvement of AgamOBP4 in insecticide recognition.
- Recognition of food-associated fragrances and environmental contaminants highlights the potential for the development of AgamOBP4-based biosensors for food quality and environmental monitoring.
- Behavioral bioassays have identified benzoyl peroxide as a potent repellent against the Asian tiger mosquito *Aedes albopictus*.
- AI-based protein-ligand modeling provided insight into binding-mode, affinity, and bioactivity relationships

## Introduction

Odorant-binding proteins (OBPs) are small, soluble proteins that play a central role in insect chemoreception. Since their initial discovery (Vogt & Riddiford, 1981) OBPs have been recognized as key contributors to insect behaviors, such as host recognition, mate selection, oviposition, and feeding, functioning as molecular carriers of odors. They bind, solubilize, and transport hydrophobic odor molecules through the aqueous sensillar lymph to the olfactory receptors (ORs) located on sensory dendrites, while also protecting them from degradation (Abendroth, Moural et al., 2023, Brito, Moreira et al., 2016, Sun, Xiao et al., 2018). By acting as the primary filters for odor selection and discrimination, OBPs enhance both the selectivity and sensitivity of the insect olfactory system in complex chemical environments (Fan, Francis et al., 2011, Leal, 2003). Furthermore, they have also been proposed to be involved in insect resistance mechanisms by sequestering or detoxifying insecticide molecules and plant-derived toxins (Deng, Xu et al., 2025, Pu & Chung, 2024, Wang, Shentu et al., 2025). Due to their functional versatility, OBPs have emerged as promising molecular targets for manipulating insect behavior in pest control strategies (Liggri, Perez-Garrido et al., 2023a, Pelosi, Iovinella et al., 2018, Rana, Sharma et al., 2024, Thireou, Kythreoti et al., 2018, Venthur & Zhou, 2018).

Computational approaches have been increasingly applied to identify bioactive molecules by exploiting known insect OBP structures. Moreover, the integration of ligand-based virtual screening with OBP structure-based design has enabled the identification of novel bioinspired repellents with enhanced efficacy, providing a powerful strategy for the development of next-generation mosquito control agents (Liggri et al., 2023a).

Although OBP amino acid sequences often show low identity across and within insect species, these proteins generally maintain a conserved structural scaffold (Manoharan, Ng Fuk Chong et al., 2013). Classical OBPs are characterized by six conserved cysteine residues that form three disulfide bonds, stabilizing an α-helical structure (Pelosi et al., 2018, Zhou, 2010). Despite this conserved architecture, OBPs display substantial structural variability, including differences in the length, orientation, and tilt of their α-helices, as well as in the arrangement of connecting loops and terminal regions. Such variations result in diversity in the size, shape, and chemical environment of their internal hydrophobic binding cavities, features closely related to the range and specificity of their ligands (Brito et al., 2016, Liggri, Tsitsanou et al., 2023b, Tegoni, Campanacci et al., 2004, Zheng, Li et al., 2015). Moreover, ligand binding to OBPs further induces structural rearrangements, such as the reorientation of helices, loops, or side chains lining the binding cavity, facilitating ligand accommodation and optimizing protein–ligand interactions, thereby enhancing both ligand specificity and affinity (Christodoulou, Stamati et al., 2026, Gonzalez, Rihani et al., 2020, Lagarde, Spinelli et al., 2011, Spinelli, Lagarde et al., 2012, Ziemba, Murphy et al., 2013).

In some OBPs, large internal cavities have been found to contain multiple binding sites, allowing two molecules of the same ligand or two distinct compounds to bind simultaneously. In the AgamOBP1–Icaridin crystal structure, Icaridin was observed to be bound to the DEET-binding site in two distinct orientations, as well as to a novel secondary site near the protein’s C-terminal region (Drakou, Tsitsanou et al., 2017). Similarly, the plant-derived compound Carvacrol occupied two binding sites of different specificities within the large cavity of AgamOBP5 (Liggri et al., 2023b). In the AgamOBP9 complex with n-butyl cinnamate, the ligand binds to the primary site located deep within the internal cavity, whereas an additional MPD molecule derived from the crystallization solution is accommodated at a secondary site near the cavity opening, serving as a surrogate ligand (Christodoulou et al., 2026).

In addition to ligand-induced structural alterations, changes in pH have also been shown to induce conformational changes in OBPs, influencing ligand binding and release processes (Briand, Nespoulous et al., 2001, Han, Zhang et al., 2014, Leal, Ishida et al., 2009, Mam, Tsitsanou et al., 2023, Manoharan, Fuchs et al., 2014, Sandler, Nikonova et al., 2000, Yin, Choo et al., 2015, Zheng, Yang et al., 2022). Such changes may occur along the olfactory pathway, from the pore wall of the antenna sensilla, which exhibits a net negative charge that lowers the local pH, to the sensillum lymph (pH ∼6.5) and further toward the vicinity of the ORs (pH ∼4.5)(Damberger, Michel et al., 2013, Keil, 1984).

The *Anopheles gambiae* AgamOBP4 protein, a member of the classic OBP subfamily, is one of the ten AgamOBPs proposed to be involved in host-seeking behavior. It is predominantly expressed in the antennae of adult female mosquitoes and is upregulated before a blood meal, suggesting its role in detecting host odors (Biessmann, Walter et al., 2002). Therefore, it has been highlighted as a potential target for discovering novel disruptors of mosquito-host interactions.

AgamOBP4 undergoes pH-dependent conformational changes. At low pH, increased flexibility of its N-terminal tail and rearrangements of helices α4 and α5 along with their connecting loop promote the enlargement of the opening to its binding cavity, potentially facilitating ligand entry. As the pH reaches neutrality, AgamOBP4 appears to exhibit increased helicity and stability, enabling the binding of indole, a human-host and oviposition attractant, with enhanced affinity (Mam et al., 2023). This further promotes the creation of a binding site for AgamOBP1 and subsequently leads to the formation of AgamOBP4-AgamOBP1 heterodimers (Andronopoulou, Labropoulou et al., 2006, Davrazou, Dong et al., 2011), with binding specificity distinct from those of the individual proteins (Qiao, He et al., 2010). Given that AgamOBP1 is a requisite protein for mosquito antenna responses to indole (Biessmann, Andronopoulou et al., 2010, Pelletier, Guidolin et al., 2010), and that AgamOBP4 has been established as an indole carrier, interference with AgamOBP4 function may affect heterodimer formation and consequently may play a crucial role in indole transport to the ORs (Andronopoulou et al., 2006, Mam et al., 2023). This hypothesis suggests an additional mechanism through which AgamOBP4-targeting ligands could disrupt mosquito host-seeking and oviposition behaviors.

The crystal structures of AgamOBP4 in the apo state (PDBids: 8C68, 8C6E) at both tested pH values (pH 4.6 and 8.5)(Mam et al., 2023) revealed a spacious hydrophobic binding cavity occupied by one indole molecule in the AgamOBP4–indole complex at pH 6.85 (PDBid: 3Q8I) (Davrazou et al., 2011). However, the large volume of this binding cavity may suggest that AgamOBP4 can accommodate bulky ligands or potentially more than one molecule simultaneously, as has been observed in AgamOBP1 and AgamOBP5 (Drakou et al., 2017, Liggri et al., 2023b). Structural comparison of the apo-AgamOBP4 at acidic (PDBid: 8C68), near-neutral (PDBid: 8C6G) and alkaline pH (PDBid: 8C6E), and with its indole-bound form at pH 6.85 (PDBid: 3Q8I), revealed that the most pronounced structural differences are located around the entrance to the binding cavity, whereas the bottom of the binding cavity remains largely conserved.

A comprehensive understanding of the characteristics of the AgamOBP4 binding cavity, including the presence of multiple binding sites, key amino acid residues involved in ligand recognition, critical protein–ligand interactions, and conformational changes associated with ligand binding and release, is crucial for structure-based strategies aimed at the discovery and design of novel agents capable of disrupting mosquito–host interactions.

Herein, we present the novel crystal structures of AgamOBP4, one in complex with the fluorescent probe 1-NPN and the other with two indole molecules bound simultaneously within the binding cavity. Together, these structures reveal the presence of two aromatic recognition subsites within the ligand-binding pocket. These structures served as the basis for a structure-based virtual screening against a curated library generated through the volatility-based filtering of chemically diverse compounds, including approved drugs (DrugBank), food ingredients (FooDB), Active Pharmaceutical Ingredients (APIs), drug-like small molecules and building blocks (BioNet), as well as natural volatiles reported in the literature. Docking analyses and subsequent evaluation proposed 23 compounds. These candidates were further assessed using *in vitro* fluorescence competitive binding assays to evaluate their binding affinities, and seven of them were subsequently subjected to behavioral assays to determine their repellent activity. Finally, an AI-based modeling approach was applied to these experimentally characterized compounds to investigate their binding modes to AgamOBP4, providing structural insights into the molecular determinants governing ligand recognition and suggesting structure-based principles that underline the relationship between binding mode, binding affinity, and biological activity.

## Materials and Methods

### Bacterial expression and purification of recombinant AgamOBP4 protein

*Anopheles gambiae* recombinant AgamOBP4 protein was expressed and purified according to previously published protocols (Mam et al., 2023). The highly purified protein was delipidated as described by Oldman et al. (Oldman et al., 2001) and concentrated to 15 mg/mL in 150 mM sodium acetate, pH 5, for use in fluorescence and crystallization experiments.

### Crystallization and Data Collection

Crystals of AgamOBP4 at pH 4.6 grew at 20°C using the sitting drop vapor diffusion method from a solution containing 15 mg/mL protein, 0.1 M sodium acetate, pH 4.6, and 2.0 M ammonium sulphate (Mam et al., 2023). Co-crystals of AgamOBP4 with 1-NPN were obtained under the same conditions in the presence of 2.5 mM 1-NPN (1% methanol) in the precipitant solution. Preformed AgamOBP4 crystals were soaked in mother liquor containing 25 mM Indole (10% DMSO) for 10 min. Prior to data collection, all crystals were transferred to their respective crystallization solution supplemented with 15% (v/v) glycerol as a cryoprotectant and flash-frozen at 100 K. Diffraction data for the AgamOBP4–1-NPN complex were collected from a single crystal at the X13 beamline at EMBL Hamburg Outstation, Germany (λ=1.08 Å). X-ray diffraction data for the AgamOBP4–Indole complex were collected using an Oxford Diffraction microfocus sealed-tube X-ray source with Cu Kα radiation (λ = 1.5418 Å) at NHRF, Greece.

### Structure Determination

Integration and data reduction were carried out using either CrysAlisPro (AgamOBP4-2xIndole) or MOSFLM (AgamOBP4-1-NPN) (Leslie, 1992) and SCALA of the CCP4 suite (COLLABORATIVE COMPUTATIONAL PROJECT, 1994) and intensities were converted to amplitudes with TRUNCATE (French & Wilson, 1978). Initial phases were determined by molecular replacement with MOLREP (Vagin & Teplyakov, 1997) using the AgamOBP4 structure at pH 4.6 (PDBid: 8C68) (Mam et al., 2023) as a search model. Models were further improved through iterative cycles of manual building in COOT (Emsley & Cowtan, 2004) and refinement using the maximum likelihood target function as implemented in the program REFMAC (Murshudov, Vagin et al., 1997). Water molecules were added to unidentified *F*_o_–*F*_c_ map peaks exceeding 1.0*σ* in COOT and were manually inspected.

Furthermore, acetate (ACT) and sulfate (SO4) ions, glycerol, and ligand molecules were modelled into the difference electron density maps and subsequently included in the refinement cycles. Details of data processing and refinement statistics are summarized in **Table 1**.

**Table 1.**
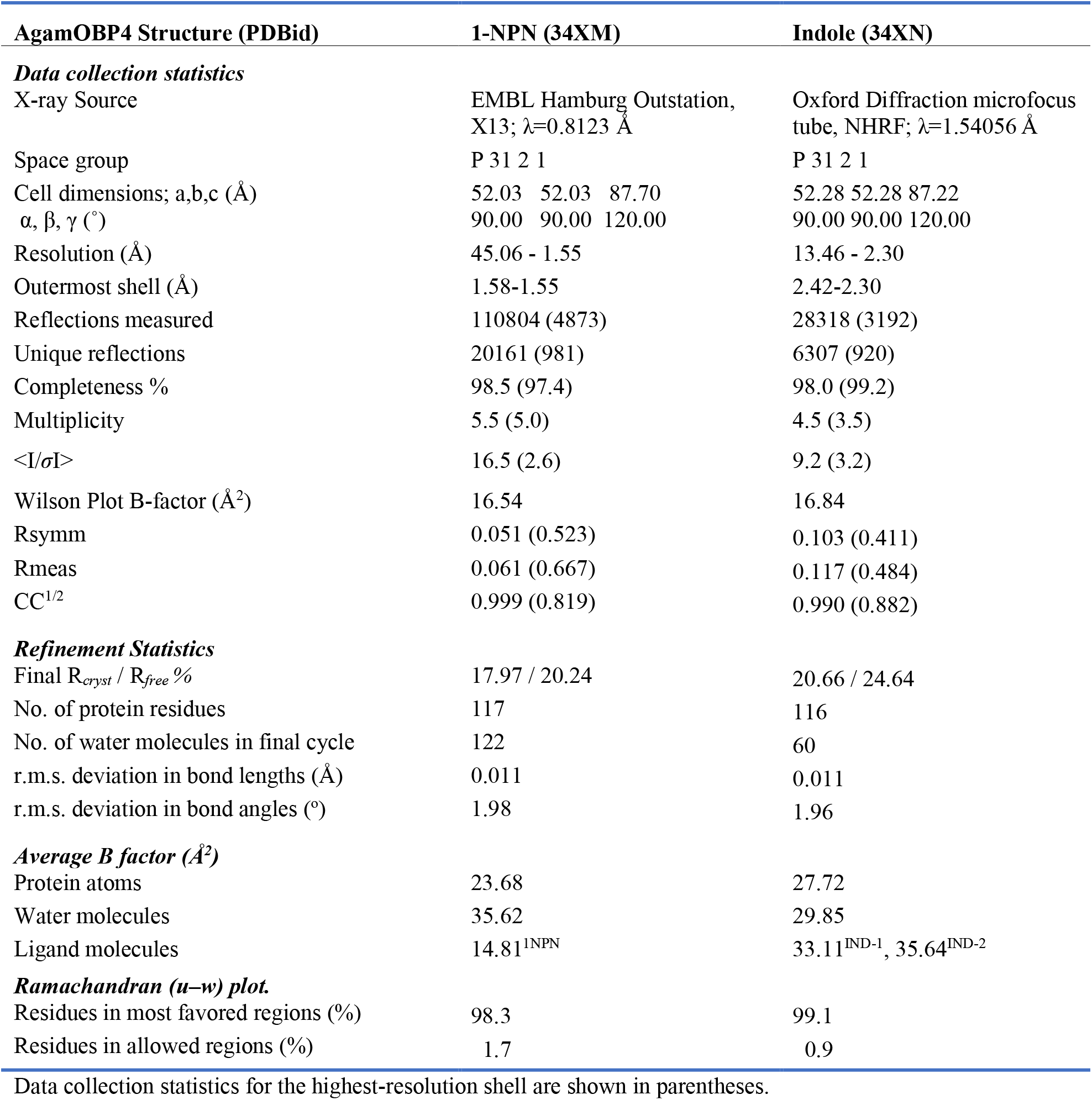
Statistics of data collection, processing and refinement of the AgamOBP4 complex structures.

| AgamOBP4 Structure (PDBid) | 1-NPN (34XM) | Indole (34XN) |
| --- | --- | --- |
| ***Data collection statistics*** |  |  |
| X-ray Source | EMBL Hamburg Outstation,<br>X13; $\lambda=0.8123$ Å | Oxford Diffraction microfocus<br>tube, NHRF; $\lambda=1.54056$ Å |
| Space group | P 31 2 1 | P 31 2 1 |
| Cell dimensions; a,b,c (Å) | 52.03 52.03 87.70 | 52.28 52.28 87.22 |
| $\alpha, \beta, \gamma$ (°) | 90.00 90.00 120.00 | 90.00 90.00 120.00 |
| Resolution (Å) | 45.06 - 1.55 | 13.46 - 2.30 |
| Outermost shell (Å) | 1.58-1.55 | 2.42-2.30 |
| Reflections measured | 110804 (4873) | 28318 (3192) |
| Unique reflections | 20161 (981) | 6307 (920) |
| Completeness % | 98.5 (97.4) | 98.0 (99.2) |
| Multiplicity | 5.5 (5.0) | 4.5 (3.5) |
| $\langle I/\sigma I \rangle$ | 16.5 (2.6) | 9.2 (3.2) |
| Wilson Plot B-factor (Å<sup>2</sup>) | 16.54 | 16.84 |
| Rsymm | 0.051 (0.523) | 0.103 (0.411) |
| Rmeas | 0.061 (0.667) | 0.117 (0.484) |
| CC<sup>1/2</sup> | 0.999 (0.819) | 0.990 (0.882) |
| ***Refinement Statistics*** |  |  |
| Final $R_{\text{cryst}} / R_{\text{free}}$ % | 17.97 / 20.24 | 20.66 / 24.64 |
| No. of protein residues | 117 | 116 |
| No. of water molecules in final cycle | 122 | 60 |
| r.m.s. deviation in bond lengths (Å) | 0.011 | 0.011 |
| r.m.s. deviation in bond angles (°) | 1.98 | 1.96 |
| ***Average B factor (Å<sup>2</sup>)*** |  |  |
| Protein atoms | 23.68 | 27.72 |
| Water molecules | 35.62 | 29.85 |
| Ligand molecules | 14.81<sup>1NPN</sup> | 33.11<sup>IND-1</sup>, 35.64<sup>IND-2</sup> |
| ***Ramachandran (u-w) plot.*** |  |  |
| Residues in most favored regions (%) | 98.3 | 99.1 |
| Residues in allowed regions (%) | 1.7 | 0.9 |

Data collection statistics for the highest-resolution shell are shown in parentheses.

The final structures quality was evaluated using PDB-REDO (Joosten, Long et al., 2014) and wwPDB Validation server (https://validate.wwpdb.org). Protein–ligand interactions within a 4.0 Å cutoff distance, as well as hydrogen bonds, were identified using CONTACT from the CCP4 suite and HBPLUS (McDonald & Thornton, 1994). 2D diagrams of ligand– protein interactions were generated with BIOVIA Discovery Studio 2020 Visualizer. RMSD plot was created with the MultiSeq module (Roberts, Eargle et al., 2006) of the VMD suite (Humphrey, Dalke et al., 1996). PyMOL (DeLano, 2002) was used to create all 3D structural figures from the molecular structure data. The atomic coordinates of AgamOBP4 complexed with 1-NPN, and indole have been deposited in the RCSB Protein Data Bank (http://www.rcsb.org/pdb) and are available under the accession codes **34XM** and **34XN,** respectively.

### Computational Methods

#### Crystal Structure Preparation for Docking

Three AgamOBP4 crystal structures were provided for computational analysis:

- 3Q8I.pdb / Crystal structure of *Anopheles gambiae* odorant binding protein 4 in complex with 1 indole molecule from RCSB (Davrazou et al., 2011)
- OBP4-1NPN_4.6.pdb / OBP4 with 1NPN at pH 4.6 (this study; PDBid **34XM**)
- OBP4-Indole_4.6.pdb / OBP4 with 2 indole molecules at pH 4.6 (this study; PDBid **34XN**)

All structures were visually inspected and processed using PyMOL. Co-crystallized ligands (1-NPN and/or indole) were identified and their coordinates recorded for reference. Subsequently, ligands and non-protein heteroatoms were removed, including water molecules and crystallization components (H₂O, PO₄⁻, SO₄²⁻, acetate, and glycerol). Cleaned protein structures were saved in PDB format. The co-crystallized ligand coordinates were kept as the center of the docking box.

Protein preparation was performed using AutoDockTools (Sanner, 1999) (included in MGLTools v1.5.7p1). Missing hydrogens were automatically added, AutoDock4 atom types were assigned, and Gasteiger partial charges were computed. Final receptor files were saved in PDBQT format.

#### Database Selection and Filtration

Compound libraries were collected from:

- Publicly available databases:

- DrugBank (version 5.1.4, 11,945 molecules) - curated database of approved and investigational drugs (Wishart, Feunang et al., 2018).
- FooDB (version pre-1.0, 70,229 molecules) - database of natural food and related compounds (www.foodb.ca).
- KeyOrganics BioNet (2024 release, 308,453 molecules) - commercially available small bioactive molecules (www.keyorganics.net). Among these, ten entries correspond to synthetic compounds displaying shape and chemical similarity to 1-NPN. These compounds were previously identified by ligand-based virtual screening of the KeyOrganics BioNet database (Complete Core Scaffolds, 2011 release; 42,755 molecules) (Thireou et al., 2018). These entries are indicated with the prefix “npnh-” in the merged BioNet library.
- Previously identified candidate sets, for a total of 337 molecules: Natural volatiles previously screened for the identification of antagonists of the *Anopheles gambiae* odorant receptor co-receptor (Kythreoti, Thireou et al., 2024)

#### Volatility filtering

Since *in vivo* the compounds would need to be volatile enough to reach mosquitoes’ olfactory system, we decided to discard molecules that do not fit typical volatility values. All compounds were saved in SMILES format to calculate vapor pressure values (mm Hg at 25 °C) using MPBPWIN v1.43 from the EPISUITE v4.11 package (US-EPA, 2026). The EPISUITE package was executed under Linux using the Wine compatibility layer (https://www.winehq.org/), which enables the execution of Windows applications on POSIX-compliant operating systems without full system emulation. The reported vapor pressure corresponded to the mean estimation derived from the Antoine and Grain methods (Meylan, Howard et al., 1996). The set of compounds was filtered to retain molecules with predicted vapor pressures between 7 × 10⁻⁷ and 1 × 10⁻¹ mmHg, consistent with the expected physicochemical range of airborne semiochemicals. Filtering, sorting, and property merging (including SMILES strings and calculated descriptors) were performed using Python and pandas.

Of 390,964 input molecules, 139,133 were retained: 133,331 from BioNet, 4053 from FooDB, 1597 from DrugBank and 152 from the collaborator sets.

#### Ligand Preparation

Filtered compounds were converted from SMILES to 3D structures using molconvert from ChemAxon (http://www.chemaxon.com). The following options were applied: 3D structure generation using the Fine Build algorithm for high-quality coordinate construction (Murshudov et al.), geometry optimization using the MMFF94 force field (Halgren, 1996) and strict convergence criterion to ensure tight gradient thresholds and improved structural refinement.

The resulting ligand files were processed with AutoDockTools’ script prepare_ligand4.py and saved to PDBQT format.

#### Docking Calculations

Virtual screening was performed using AutoDock Vina v1.1.2 (Trott & Olson, 2010) through the MetaScreener suite (https://github.com/bio-hpc/metascreener), employing default values. All compounds were screened against each structure, generating poses and docking scores which were later merged into a single table for comparison.

#### Ranking and Selection

Docking score tables generated by MetaScreener were consolidated and merged with previously calculated physicochemical properties (vapor pressure, molecular weight, and related descriptors). 2D interaction diagrams were generated with PoseView 1.1.2 (BioSolveIT / ZBH, University of Hamburg) (Stierand, Maass et al., 2006). Because docking scoring functions tend to favor larger compounds due to increased contact surface area, a metric consisting on dividing docking score by molecular weight was computed to account for size-related bias. This adjustment is consistent with previously reported observations in structure-based virtual screening studies and scoring validation (Pan, Huang et al., 2003).

#### AI-based modelling of ligand binding

The seven experimentally validated ligands listed in **Table 4** were modeled in complex with the receptor using Protenix. Protein–ligand complex structures were predicted using the AI-based deep-learning model protenix_base_constraint_v0.5.0 through the Tamarind Bio platform (https://app.tamarind.bio/protenix). Protenix is an open-source structure-prediction framework based on the AlphaFold 3 architecture (Abramson, Adler et al., 2024), which employs a diffusion-based generative approach to predict three-dimensional biomolecular structures from sequence and chemical information (ByteDance, Chen et al., 2025). For the present study, predictions were generated from the AgamOBP4 amino-acid sequence (Q8T6R7; residues 26-150; Supplementary information) and the SMILES representation of each ligand. The experimentally determined ligand-binding site of AgamOBP4, defined from the crystallographic structure of AgamOBP4-1-NPN (PDBid **34XM**) (residues 52, 53, 56, 58, 65, 67, 70, 73, 93, 94, 107, 110, 111, 114, 124), was incorporated as a soft pocket constraint, with a maximum ligand–binding-site residue distance of 8 Å. Multiple Sequence Alignment (MSA) providing evolutionary information to support protein structure prediction and Training-Free Guidance (TFG) was enabled during inference to improve plausibility of the predicted ligand poses, respectively. The resulting protein–ligand complexes were evaluated by superimposing the predicted AgamOBP4 structures onto the crystallographic structure using the SUPERPOSE program from the CCP4 suite. The predicted ligand positions were subsequently compared with the crystallographically determined 1-NPN and Indole binding modes and the associated protein–ligand interactions. The RMSDs of the predicted 1-NPN and indole poses relative to their respective crystallographic poses were calculated with RDKit: Open-source cheminformatics. https://www.rdkit.org, version 2026.03.5 (https://doi.org/10.5281/zenodo.591637).

### Fluorescent measurements

The binding affinities of the selected compounds for AgamOBP4 were indirectly determined from the displacement of the fluorescent probe 1-NPN (N-phenyl-1-naphthylamine) by the ligand through a fluorescence competitive binding assay. The dissociation constant (Kd) of 1-NPN for AgamOBP4 was obtained by titrating increasing concentrations of 1-NPN (0, 0.1, 0.2, 0.3, 0.5, 1, 2, 3, 5, 7, 9, 15, 20, 30, 40 μM) in a solution containing 2 μΜ protein in 20 mM Tris-HCl (pH 8) and 100 mM NaCl.

The affinity of the docking-hit compounds for AgamOBP4 was initially assessed using a one-point fluorescence assay under the same buffer conditions, employing 2 μM protein and 2 μM 1-NPN, with each compound tested at a single concentration of 30 μM. Ligand binding affinity was expressed as the percentage of fluorescence intensity relative to the fluorescence of the AgamOBP4–1-NPN complex in the absence of ligand, which corresponds to 100% probe bound to the protein.

To determine the dissociation constants (Ki) for compounds that induced more than 55% fluorescence reduction in the one-point screening assays, the protein was assayed under identical experimental conditions in the presence of increasing ligand concentrations ranging from 0 to 60 μM. The assays were performed in black 96-well plates (Greiner Bio-One), in a final reaction volume of 300 μL per well. Fresh aliquots of 1-NPN and the various ligands were prepared in 100% HPLC-grade DMSO, ensuring that the final concentration of the organic solvent in the assay remained below 2.7% to avoid protein denaturation.

The fluorescent probe was excited at a wavelength of 337 nm, and the emission spectra were recorded from 360 to 500 nm on a Varioskan Flash fluorimeter plate reader (Thermo Scientific) at a stable temperature of 25 °C.

To account for potential interference with the fluorescent probe, control samples containing 1-NPN and ligand in the absence of protein were analyzed in parallel. The resulting background fluorescence was subtracted from the corresponding experimental measurements before further data processing. All experiments were performed in triplicate to ensure reproducibility. The dissociation constant (Kd) of 1-NPN and the Ki’s of the ligands were calculated by the nonlinear regression analysis of GraphPad Prism 8 (www.graphpad.com).

The Kd of 1-NPN for AgamOBP4 was determined to be 6.29 μM, from a saturation curve generated by fitting the fluorescence data to the one-site binding model equation **(Supplementary Figure S1)**:

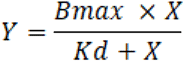

where Bmax is the maximum binding capacity of the protein, X corresponds to the total ligand concentration in the assay, and Kd is the equilibrium dissociation constant.

The % fluorescence decrease data were fitted to an “[Inhibitor] vs response” model using the following equation:

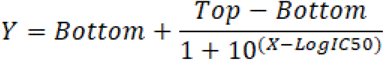

where Top and Bottom represent the upper and lower plateaus of the fluorescence responses, respectively, and IC50 the ligand concentration that corresponds to a % fluorescence intensity which is halfway between these two values. Because the lower asymptotes (Bottom plateaus) of the displacement curves did not reach zero and exhibited variation among the tested ligands, the IC50int value was further calculated. IC50int is defined as the ligand concentration required to reduce the initial fluorescence by 50% and is determined by interpolating the displacement curves using the built-in functions of GraphPad Prism. Subsequently, the dissociation constant (*K*i) of each ligand was calculated using the Cheng-Prusoff equation for competitive conditions:

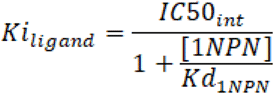

In this equation [1-NPN] represents the concentration of the probe in the assay (2 μM), and Kd_1NPN_ is its calculated dissociation constant (6.29 μM; **Figure S1**).

### Rearing of *Aedes albopictus* in the laboratory

A laboratory colony of Aedes albopictus was maintained under controlled environmental conditions (28 °C constant temperature, 80% relative humidity, and a 16:8 h light:dark photoperiod) at the Insectary of the Benaki Phytopathological Institute, Kifissia, Greece (Chazapi, Kritsi et al., 2025). Larvae were fed ad libitum with fish food pellets (Goldfish Sticks, Tetra) until pupation. Pupae were then transferred to BugDorm^®^ cages (32 × 32 × 32 cm) covered with a 32-mesh screen. Adult mosquitoes were maintained on a 10% sucrose solution provided ad libitum. For colony maintenance, female mosquitoes were blood-fed using a Hemotek membrane feeding system (FAO/IAEA) at the BPI laboratory, an authorized facility for the use of animal by-products (registration no. AK18Y3EL25XP, Region of Attica).

### Repellent activity bioassay

The repellent activity of the selected compounds was assessed using the human landing count method, following previously published protocols (Chazapi et al., 2025, Giatropoulos, Papachristos et al., 2012). Dichloromethane (DCM) was used as control and each treatment was tested in eight replicates using four human volunteers in rotation. Each replicate consisted of a 5-min exposure period.

### Determination of repellency index

The landing numbers were converted to repellency index (RI±SE) using the following equation

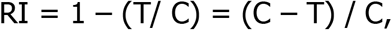

where C is the number of landings in the control and T the number of landings in the compound treatment.

### Statistical analysis

Landing numbers were converted to repellency indices (RI) using the following equation: RI = [1 - T/C] x 100, where C is the number of landings in the control and T is the number of landings in the treatment. Data concerning the repellency indices were analyzed using the Kruskal–Wallis test (Kruskal & Wallis, 1952).. When significant differences were detected, Mann–Whitney U tests were carried out for pair-wise comparison with a Bonferroni correction for adjustment of p-values (Mann & Whitney, 1947).

## Results

### Overview of structural features of AgamOBP4-ligand crystal complexes

Structure-based approaches for identifying potential protein binders rely on experimentally determined protein structures, particularly protein-ligand complexes, which define the location, geometry, and physicochemical characteristics of the ligand binding sites, commonly situated within internal cavities. In the case of AgamOBP4, previously reported structures at three different pH conditions (Mam et al., 2023) revealed a relatively large and preformed binding cavity, partially occupied by an indole molecule in the AgamOBP4-indole complex at pH 6.85 (PDBid: 3Q8I). This observation suggests that the AgamOBP4 cavity could potentially accommodate bulkier compounds or even multiple ligands simultaneously. To explore this possibility, 1-NPN (N-phenyl-1-naphthylamine), commonly used as a fluorescent probe in competitive binding assays, was included in co-crystallization trials, while preformed AgamOBP4 crystals were subjected to soaking with high concentrations of indole.

AgamOBP4 exhibits the characteristic features of classical OBPs, consisting of six α-helices, α1 (aa. 1-17), α2 (aa. 25-33), α3 (aa. 42-55), α4 (aa. 66-76), α5 (aa. 82-91) and α7 (aa. 102-117), stabilized by conserved disulfide bonds. The main entrance to the internal protein cavity is located between α1, α3, and α4 helices. At pH 4.6, the N-terminal helix segment of AgamOBP4 (residues 1 to 8) exhibits increased flexibility, resulting in the widening of the opening, possibly facilitating the entrance and exit of the ligand. In addition, a second, narrower opening to the cavity is also observed, located on the opposite side of the protein, between helices α4 and α5.

As both AgamOBP4–1-NPN and AgamOBP4–indole structures were obtained from crystals grown under acidic conditions (pH 4.6), no electron density was detected for residues 1–8. In particular, in the AgamOBP4–1-NPN complex, obtained via co-crystallization, the persistence of this disorder suggests that ligand binding does not influence the stability of this region. The structural integrity of the remaining regions of AgamOBP4 at low pH was further reflected in its crystallization behavior, which yielded larger and higher-quality crystals than those obtained under neutral or alkaline conditions, suitable for high-resolution structure determination. Thus, acidic pH substantially facilitated the crystallographic investigation of AgamOBP4 while preserving the overall architecture of the ligand-binding site.

Structural analysis of the AgamOBP4–1-NPN complex, solved at 1.55 Å resolution, revealed a 1-NPN molecule bound at the distal end of the internal cavity, positioned in front of the secondary opening and effectively blocking access through this entrance (**Figure 1Α**). An extra positive electron density observed in the remaining area of the cavity, which could not be accounted for by a second ligand molecule, was assigned to a glycerol molecule and an acetate ion originating from the crystallization solution. Upon binding, 1-NPN forms a direct hydrogen bond through its nitrogen atom with the sidechain oxygen of Thr70 from the α4 helix (3.02 Å). In addition, a π–hydrogen bond is also formed between the hydroxyl group of Thr58, acting as a hydrogen bond donor, and the aromatic π-system of the benzene ring of 1-NPN, which serves as the acceptor, thereby further stabilizing the ligand within the binding site (**Figure 1Β**).

**Figure 1.**
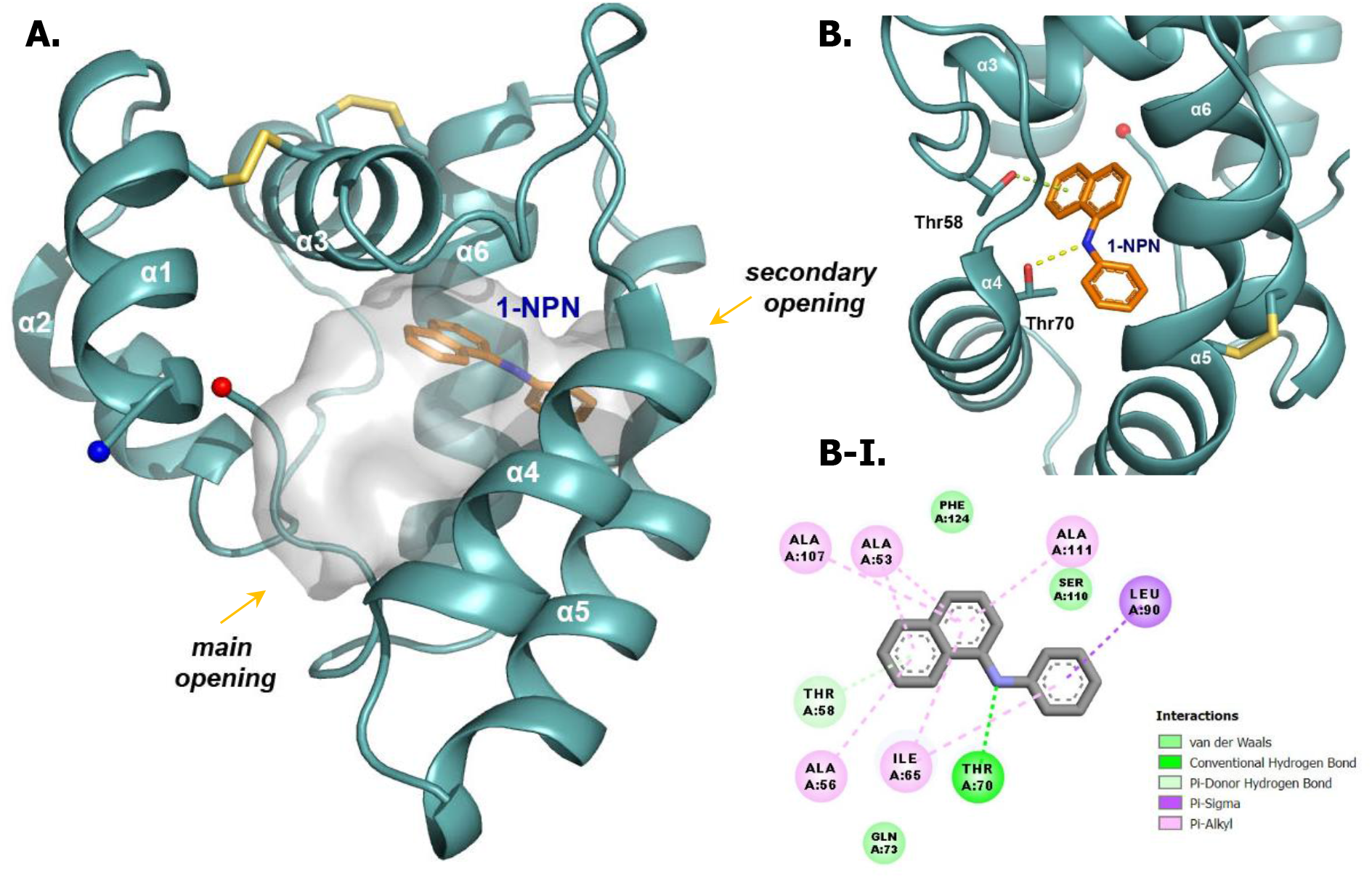
Binding of the fluorescent probe 1-NPN to AgamOBP4. **A.** Cartoon representation of the AgamOBP4 protein with 1-NPN bound at the bottom of its internal hydrophobic cavity. The main and secondary openings of the cavity are indicated. **B.** 1-NPN forms a direct hydrogen bond (yellow dashed line) with Thr70 and a π-hydrogen bond (green dashed line) to Thr58. **B-I.** 2D interaction diagram illustrating the contacts between AgamOBP4 residues and 1-NPN.

Moreover, 1-NPN participates in 54 van der Waals (vdW) contacts with residues lining the secondary opening belonging to helices α3 (Ala53), α4 (Gln73), and α6 (Ala107, Ser110, Ala111), as well as the α3-α4 connecting loop (Thr58 and Ile65), the C-terminal loop (Phe124), and the glycerol molecule (**Figure 1B-I**). Among them, π-alkyl and π-sigma hydrophobic interactions between the naphthalene and phenyl rings of 1-NPN and protein residues, although weaker than hydrogen bonds, contribute significantly to ligand stabilization.

Previous structural studies demonstrated that indole binds within the large hydrophobic cavity of AgamOBP4 at a pH of 6.85 (Davrazou et al., 2011). In this structure, indole was modelled with its NH group oriented toward the hydroxyl group of Thr70 (corresponding to Thr69 in PDB entry 3Q8I), enabling the formation of a hydrogen bond. Notably, the crystal structure of AgamOBP4 obtained by soaking preformed apo crystals at pH 4.6 with a high concentration of indole revealed the presence of two ligand molecules occupying distinct regions of the cavity (**Figure 2A**). The superposition of the two structures showed that the first indole molecule (**IND1**) occupies the same position as at pH 6.85, adopting a nearly identical planar orientation within this binding subsite. In the AgamOBP4 complex at pH 4.6, IND1 is better accommodated within the site and forms a stronger hydrogen bond through its nitrogen atom with the side-chain oxygen of Thr70 (3.1 Å), compared to the 3.6 Å distance observed at pH 6.85. Additionally, 27 vdW contacts are established with residues surrounding the subsite (Ala49, Ala53, Ala56, Ile65, Gln73, Ala107, and Phe124) and a water molecule. The π–hydrogen bond occurred in the AgamOBP4–1-NPN structure involving Thr58 is also retained, with the pyrrole ring of IND1 being the hydrogen bond acceptor (**Figure 2A and 2A-I**).

**Figure 2.**
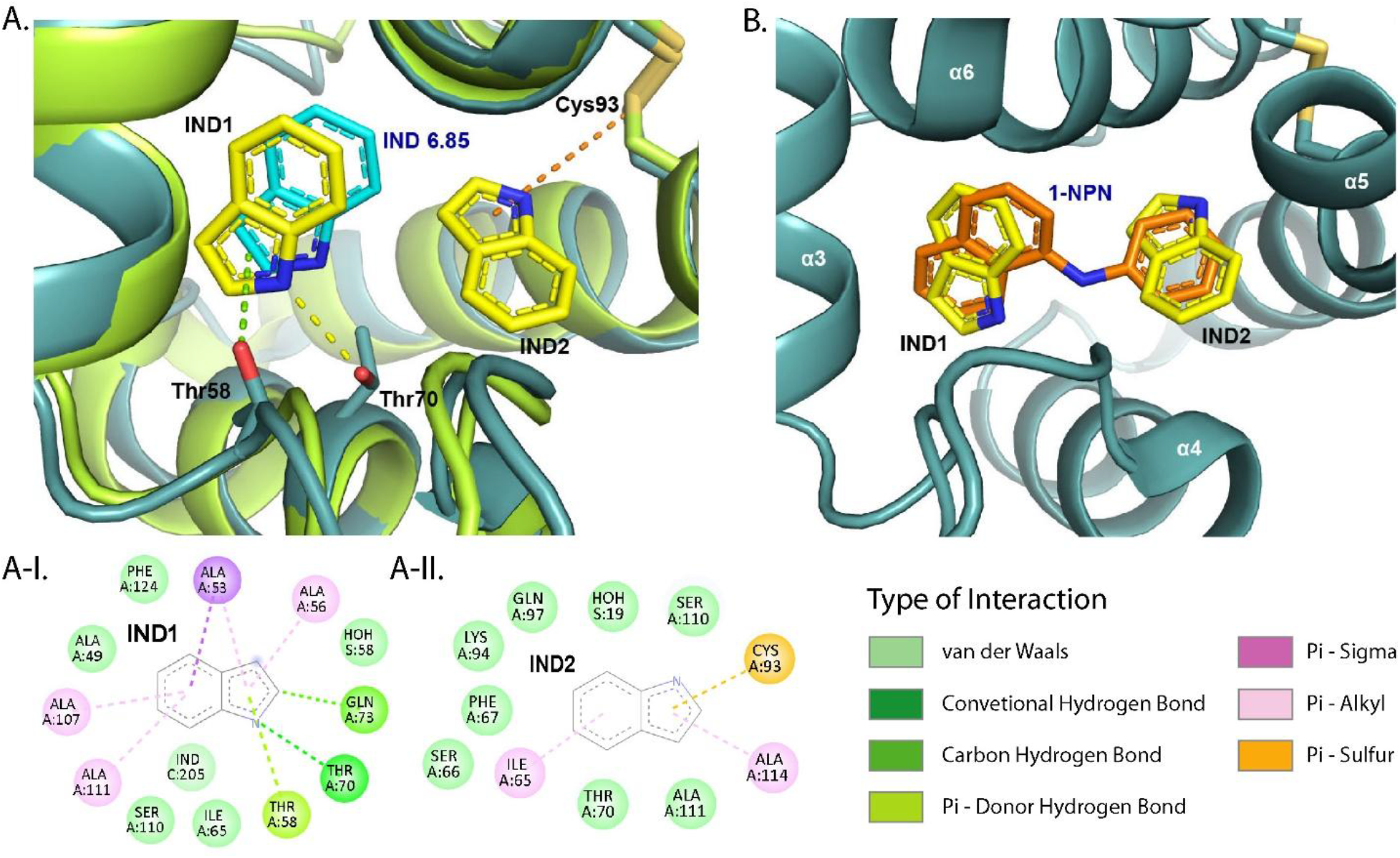
Superposition of the AgamOBP4-indole complex at pH 4.6 onto the indole-bound structure at pH 6.85 (PDBid: 3Q8I) in the binding site region. A. Two indole molecules were found to bind within the AgamOBP4 binding site, with one of them closely superimposed on the indole of the pH 6.85 structure. A shorter hydrogen bond (shown as a yellow dashed line) is formed with Thr70, while the π-hydrogen bond with Thr58 (green dashed line) observed in the 1-NPN complex is retained. The IND2 molecule participates in hydrophobic interactions with surrounding protein residues. In addition, π-sulfur contacts (brown dashed line) between its pyrrole ring and Cys93 may further stabilize its binding. A-I. and A-II. 2D diagrams of the interactions of IND1 and IND2, respectively, with AgamOBP4 residues. B. Superimposition of the two indoles (yellow) on the 1-NPN molecule (orange) at the binding site region. The indoles are colocalized with the naphthalene and phenyl rings of 1-NPN, revealing the presence of two subsites that could preferentially bind aromatic moieties.

A second indole molecule (**IND2**) binds to a neighboring subsite at the bottom of the cavity. The IND2 molecule was modelled into electron density and refined in an orientation that optimizes its interactions with the surrounding protein environment. Although it does not form any hydrogen bond, it participates in several hydrophobic contacts with surrounding amino acids and IND1 (27 vdWs), including π–sulfur interactions through its aromatic pyrrole ring with Cys93, which likely contribute to its stabilization at this site (**Figure 2A and 2A-II**).

The average root mean square deviation (RMSD) values for Cα atoms of the 1-NPN and indole (pH 4.6) complexes are 0.272 Å and 0.454 Å, respectively, compared to the apo AgamOBP4 structure at pH 4.6 (PDBid: 8C68), indicating a high degree of structural similarity (**Supplementary Figure S2**). Moderate structural deviations are observed for the first defined N-terminal helix residue Ser10 (RMSD: 1.99 Å) in the AgamOBP4–indole complex. This residue lies adjacent to the disordered N-terminal tail and is therefore expected to exhibit increased flexibility. Similarly, the C-terminal region residue Met123, which has RMSD values greater than 1.5 Å in both ligand complexes, has also been found to exhibit structural variability at different pH conditions. In addition, a slight conformational difference is observed in the segment spanning residues 58–63. Within this region, Lys61 and Lys62, located at the top of the α3–α4 connecting loop, exhibit deviations of 1.63 Å and 1.24 Å, respectively, relative to apo AgamOBP4, likely reflecting local rearrangements. Importantly, these residues are positioned far from the ligand-binding site, orienting toward the solvent.

Interestingly, superimposition of the AgamOBP4-indole structure at pH 4.6 on the AgamOBP4-1-NPN complex revealed that the two indole molecules, IND1 and IND2, closely align with the naphthalene and phenyl groups of 1-NPN, respectively (**Figure 2B**). This spatial correspondence further supports the presence of two subsites within the binding cavity, each of which exhibits a preference for aromatic ring systems. Together, these subsites appear to form a dual aromatic recognition site, capable of accommodating either multiple aromatic compounds simultaneously or larger molecules containing aromatic moieties. Furthermore, the hydrogen bond with the Thr70 sidechain appearing in both complexes is possibly significant for ligand binding to the binding site.

### Computational screening identified putative disruptors of mosquito host-seeking behavior

A virtual screening workflow was applied to identify candidate ligands targeting AgamOBP4 with potential to disrupt mosquito host-seeking behavior (**Figure 3**).

**Figure 3.**
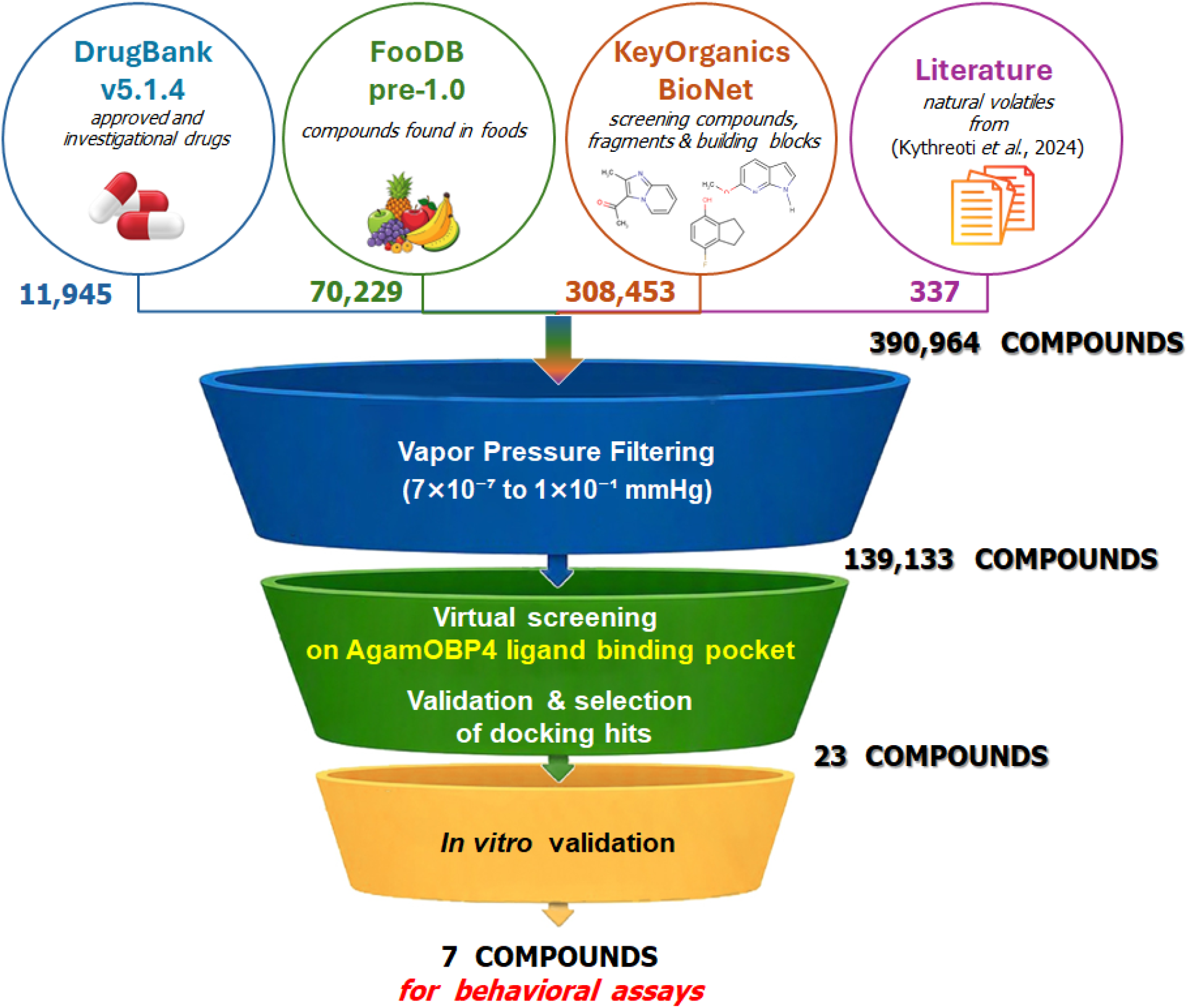
Screening protocol. Compounds from DrugBank (Wishart et al., 2018), FooDB, KeyOrganics BioNet, and a collection of natural volatiles reported by (Kythreoti et al., 2024) were filtered and screened against multiple AgamOBP4 structures. The number of compounds retained at each stage is shown.

First, the compounds collected from the source libraries (390,964 molecules) were filtered by predicted vapor pressure. Of these, 139,133 were retained. Then, docking simulations were performed across multiple AgamOBP4 crystal structures, generating binding scores and predicted ligand poses for all filtered compounds. **Supplementary Figure S3** presents the predicted binding poses of selected docking hits in the AgamOBP4 structures for which the most favorable docking scores were obtained. Notably, the docking protocol successfully identified indole among the candidate ligands, yielding moderate docking scores that are broadly consistent with its previously reported moderate affinity for AgamOBP4 (*K*d = 26.6 μM) (Mam et al., 2023). The results were subsequently analyzed to select the most promising hits. To reduce known biases toward larger molecules in docking scoring functions, scores were further adjusted by molecular weight. The table below shows a summary of the docking scores, other details, and a summary of the additional considerations for the selected compounds (**Table 2**).

**Table 2.** Compounds chosen for *in vitro* affinity evaluation at a single ligand concentration. The table lists the compound identifier (**ID**), compound number (**#**), sum of docking scores (**s-sum**) and minimum docking score (**s-min**) across all crystal structures, target structure associated with the minimum score (**best target**), predicted **volatility** in mm Hg, molecular weight (**MW**), manual pose and data evaluation score (**pev**), and manual compound evaluation score (**xev**). Compounds selected for subsequent quantitative affinity determination using fluorescence-binding assays are indicated in **bold**.

| ID | # | s-sum | s-min | best target | volatility | MW | pev | xev |
| --- | --- | --- | --- | --- | --- | --- | --- | --- |
| <sup>1</sup>FS-4526 | 1 | -70.6 | -15.3 | OBP41NPN46 | $3.1 \cdot 10^{-2}$ | 580 | 7.2 | 0 |
| FS-4985 | 2 | -49.0 | -11.2 | OBP41NPN46 | $9.5 \cdot 10^{-2}$ | 674 | 2.0 | 0 |
| <sup>2</sup>**FDB007671** | **3** | **-44.2** | **-10.7** | **OBP41NPN46** | **$1.3 \cdot 10^{-3}$** | **166** | **5.9** | **1** |
| FDB011272 | 4 | -41.2 | -10.7 | OBP41NPN46 | $1.1 \cdot 10^{-4}$ | 226 | 5.9 | 1 |
| **FDB031103** | **5** | **-41.1** | **-10.6** | **OBP41NPN46** | **$1.1 \cdot 10^{-4}$** | **226** | **5.9** | **1** |
| FDB003914 | 6 | -43.4 | -10.4 | OBP41NPN46 | $7.3 \cdot 10^{-4}$ | 226 | 5.9 | 1 |
| <sup>3</sup>kyth-7249 | 7 | -33.5 | -8.2 | OBP41NPN46 | $9.6 \cdot 10^{-2}$ | 122 | 4.1 | 0 |
| DS-9432 | 8 | -41.9 | -10.5 | OBP41NPN46 | $8.1 \cdot 10^{-4}$ | 153 | 3.2 | 0 |
| AS-10018 | 9 | -51.5 | -12.1 | OBP41NPN46 | $2.2 \cdot 10^{-4}$ | 256 | 1.3 | 1 |
| AS-58219 | 10 | -49.1 | -12.2 | OBP41NPN46 | $1.1 \cdot 10^{-4}$ | 214 | 1.3 | 1 |
| <sup>4</sup>vlat-C-12 | 11 | -22.9 | -5.0 | OBP4Indole46 | $3.4 \cdot 10^{-2}$ | 85 | 1.8 | 0 |
| **BS-51734** | **12** | **-50.3** | **-12.0** | **OBP41NPN46** | **$2.3 \cdot 10^{-4}$** | **236** | **1.3** | **1** |
| FDB016853 | 13 | -45.5 | -10.8 | OBP41NPN46 | $4.2 \cdot 10^{-4}$ | 216 | 5.9 | 1 |
| **FDB097217** | **14** | **-51.5** | **-11.1** | **3Q8I** | **$8.9 \cdot 10^{-6}$** | **318** | **6.0** | **0** |
| **FDB013700** | **15** | **-45.5** | **-10.4** | **OBP41NPN46** | **$7.5 \cdot 10^{-4}$** | **200** | **5.9** | **1** |
| kyth-5281515 | 16 | -38.6 | -8.8 | OBP41NPN46 | $2.1 \cdot 10^{-2}$ | 204 | 6.6 | 0 |
| vlat-C-73 | 17 | -37.6 | -9.0 | OBP41NPN46 | $4.6 \cdot 10^{-2}$ | 204 | 4.9 | 0 |
| <sup>5</sup>**DB05592** | **18** | **-50.5** | **-12.0** | **OBP41NPN46** | **$3.7 \cdot 10^{-6}$** | **281** | **6.0** | **0** |
| kyth-74563 | 19 | -24.6 | -5.0 | OBP41NPN46 | $6.6 \cdot 10^{-2}$ | 116 | 1.8 | 0 |
| DB09304 | 20 | -54.4 | -13.6 | OBP41NPN46 | $2.0 \cdot 10^{-6}$ | 261 | 6.2 | 0 |
| DB18178 | 21 | -53.2 | -12.1 | OBP41NPN46 | $3.7 \cdot 10^{-6}$ | 280 | 7.2 | 0 |
| **DB09096** | **22** | **-42.2** | **-10.5** | **OBP41NPN46** | **$4.2 \cdot 10^{-4}$** | **242** | **5.9** | **1** |
| kyth-798<br>(Indole) | 23 | -32.9 | -8.0 | OBP41NPN46 | $2.1 \cdot 10^{-2}$ | 117 | 3.1 | 0 |

Identifier forms are as follows, <sup>1</sup> **AA-0000**, BioNet database; <sup>2</sup> **FDB00000**, FooDB database; <sup>3</sup> **kyth-00000**, natural volatiles listed in Supplementary spreadsheet *mmc2.xlsx* of (Kythreoti et al., 2024); <sup>4</sup> **vlat-C-00**, natural volatiles reported by Kythreoti et al. (2024) and references therein (Kythreoti et al., 2024); <sup>5</sup> **DB00000**, DrugBank database.

### *In vitro* binding affinity determination of the selected docking hits for AgamOBP4

To evaluate the binding potential of the 22 predicted docking hits to AgamOBP4, an *in vitro* screening process was initially performed at a single ligand concentration by using a fluorescence-based competitive binding assay and 1-NPN as the fluorescent probe. The screening identified seven candidates that induced the highest reduction in fluorescence, with decreases exceeding 55% at a ligand concentration of 30 μM (**Figure 4**). Among these, compounds #12 and #15 exhibited direct interaction with 1-NPN, as indicated by increased fluorescence in control samples lacking protein, which prevented accurate quantification of fluorescence reduction. However, they were also included among the compounds selected for subsequent behavioral studies to assess their potential repellent activity.

**Figure 4.**
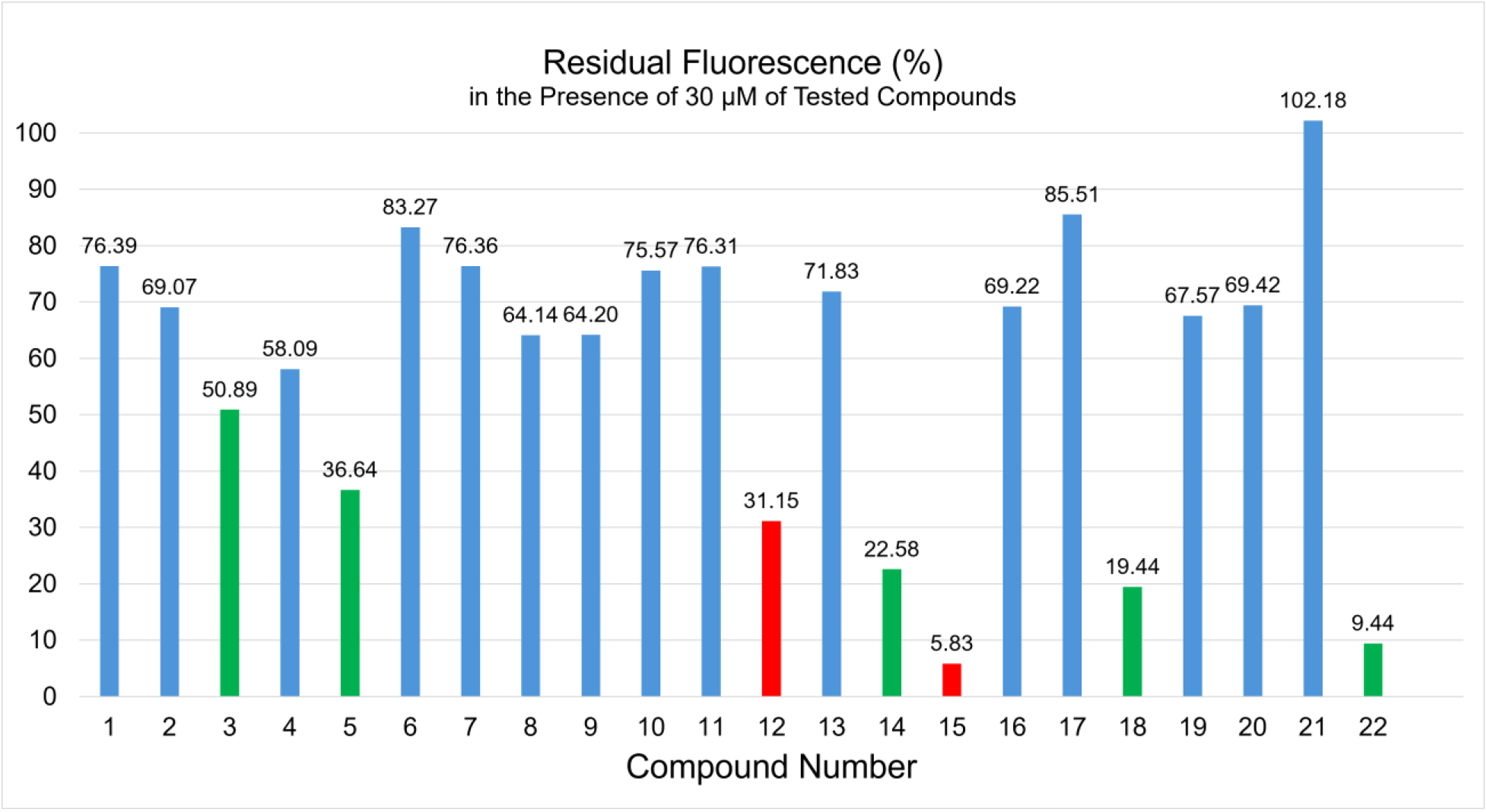
One-point screening of docking hits 1-22. The bar chart shows the percentage fluorescence intensity measured for each ligand at a final concentration of 30 μM in the presence of 2 μM 1-NPN. Results are expressed as the fluorescence intensity in the presence of the ligand relative to that of the AgamOBP4–1-NPN complex. Bars are color-coded according to compound activity: green bars indicate compounds that reduce fluorescence by more than 55%, blue bars represent compounds that induce less than 55% fluorescence reduction, and red bars denote compounds that directly interact with the 1-NPN probe.

Following the initial screening, compounds #3, #5, #14, #18, and #22, which induced the greatest reduction in fluorescence (approximately 50–90%), were selected for a detailed fluorescence binding study to determine their affinity for AgamOBP4. The titration assays were carried out under conditions of constant AgamOBP4 (2 μM) and 1-NPN (2 μM) concentrations, with the test compounds added gradually to final concentrations of up to 60 μM. Fitting the fluorescence data to a one-site saturation binding model was sufficient to describe the experimental results, indicating the presence of a single binding site on AgamOBP4 for all tested compounds (**Figure 5**). The fluorescence analysis confirmed binding of all five ligands tested, with *K*i values in the micromolar range (**Figure 5 - inset table**). However, the dissociation constant (*K*i) for compound #3 could not be determined, as the fluorescence decrease did not exceed the 50% threshold required for reliable quantification, even at the highest ligand concentration tested (60 μM). In contrast, the remaining four ligands exhibited relatively high affinity for AgamOBP4, with *K*i values ranging from 2.4 to 21.8 μM. Compound #18 was identified as the strongest binder (*K*i = 2.4 μM), followed by compound #22 (*K*i = 3.1 μM) and compound #14 (*K*i = 3.6 μM), while compound #5 displayed the weakest affinity within this group, with a *K*i value of 21.8 μM.

**Figure 5.**
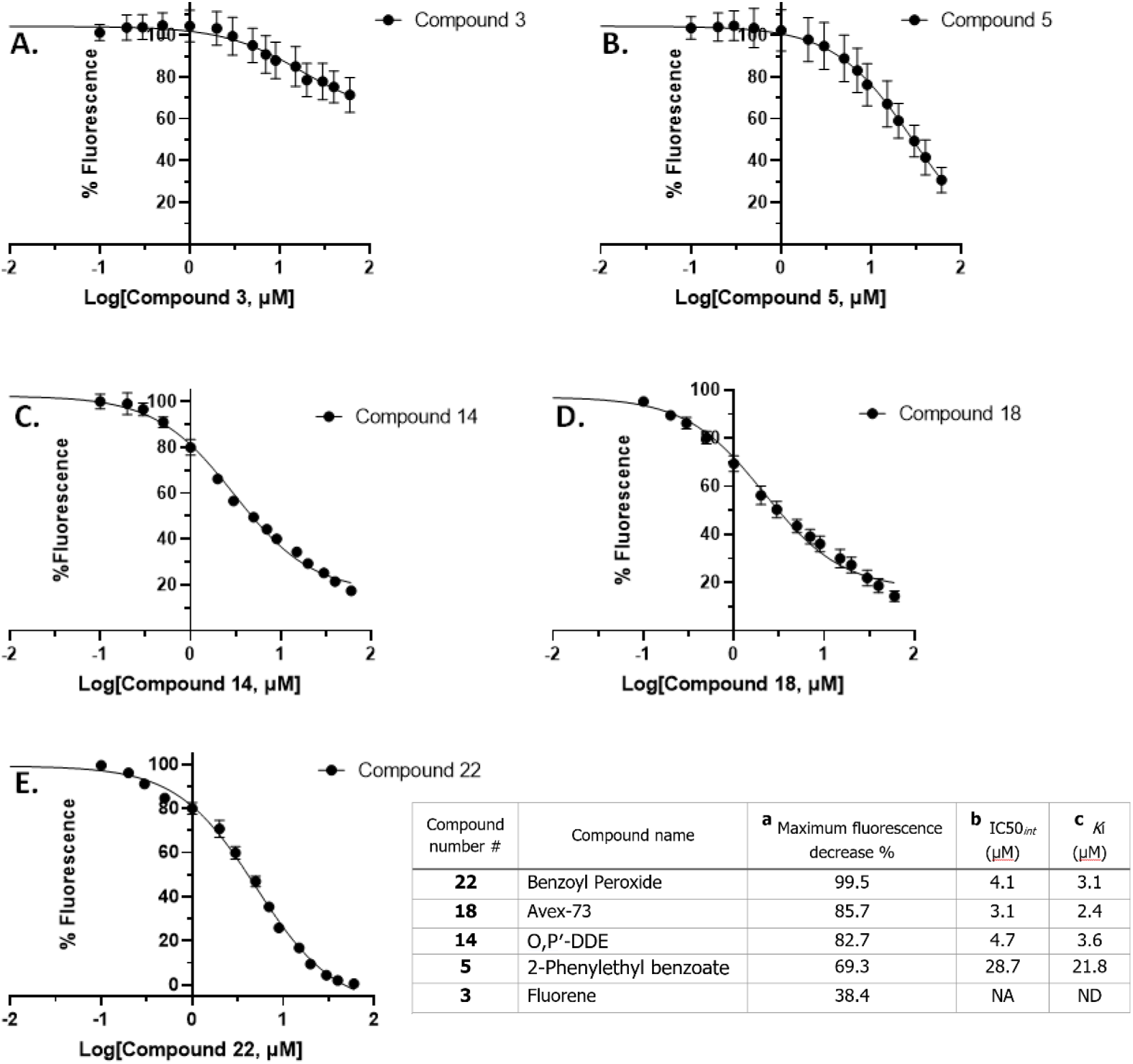
Dose–response curves illustrating the effect of ligand concentration on the displacement of the fluorescent probe 1-NPN from AgamOBP4. The plots show the percentage decrease in fluorescence intensity upon titration with increasing concentrations of compounds (**A**) #3, (**B**) #5, (**C**) #14, (**D**) #18, and (**E**) #22. (**Inset table**) Binding characteristics of the AgamOBP4 binders: **^a^** Maximum fluorescence reduction observed at the highest ligand concentration tested (60μΜ), expressed as a percentage relative to the initial fluorescence (set to 100%). **^b^** Ligand concentration required to reduce the initial fluorescence intensity by 50%. **^c^** Ligand dissociation constant (*K*i) calculated from the IC_50_int using the Cheng-Prusoff equation.

#### Effect of the selected host-seeking disruptors on Aedes albopictus mosquitoes

The four compounds characterized *in vitro* as strong AgamOBP4 binders (#5, #14, #18, and #22), together with the weak binder fluorene (#3) and the two compounds for which binding was not characterized (#12 and #15), were evaluated for repellent activity against *Aedes albopictus* mosquitoes using an arm-in-cage behavioral assay. All seven compounds were initially tested at a standard dose of 0.2 mg/cm² (**Table 3**). Using an RI >50% as the threshold for significant repellency, only benzoyl peroxide (#22) exhibited high repellent activity at this dose, with an RI of 81.8% and a statistically significant difference (group a). o,p′-DDE (#14), 2-phenylethyl benzoate (#5), fluorene (#3), and Avex-73 (#18) displayed mild repellent activity (RI = 30–50%), with Avex-73 showing the highest activity in this group (RI = 49.5%) and statistically significant differences (group a–b). Compounds #12 and #15 exhibited low repellent activity, with RIs of approximately 25%.

**Table 3.** Binding affinity and repellency of tested compounds. Dissociation constants (***K*i**) for binding to AgamOBP4 were determined using a fluorescence competition assay. Mean Repellency Index (**%RI**) was determined against *Aedes albopictus* mosquitoes using a “hand-in-cage” behavioral assay. Predicted vapor pressures (**VP**) are also listed.

| Comp. number (#) | Name | Structure | VP (mmHg) | K<sub>i</sub> (μM) | <sup>1</sup>RI%± SE<br>*Dose 0.2 mg/cm<sup>2</sup> | RI%± SE<br>*Dose 0.4 mg/cm<sup>2</sup> |
| --- | --- | --- | --- | --- | --- | --- |
| 3 | Fluorene |  | 1.3 x 10<sup>-3</sup> | NA | 37.9 ± 6.2<sup>b</sup> | 79.2±5.0<sup>a</sup> |
| 5 | Benzoic acid 2-phenylethyl ester<br>(2-Phenylethyl benzoate) |  | 1.1 x 10<sup>-4</sup> | 21.8 | 37.0±10.9<sup>b</sup> | 57.5±11.0<sup>ab</sup> |
| 12 | (4-Methylpent-1-ene-2,4-diyl) dibenzene |  | 2.3 x 10<sup>-4</sup> | ND | 26.4±16.8<sup>b</sup> | 36.6±14.9<sup>b</sup> |
| 14 | Benzene, 1-chloro-2-[2,2 dichloro-1-(4-chlorophenyl)ethenyl]<br>(o,p'-DDE) |  | 8.9 x 10<sup>-6</sup> | 3.6 | 34.0±8.9<sup>b</sup> | 40.6±6.3<sup>b</sup> |
| 15 | 2-isobutoxynaphthalene |  | 7.5 x 10<sup>-4</sup> | ND | 23.7±18.1<sup>b</sup> | 50.5±4.6<sup>b</sup> |
| 18 | Blarcamesine<br>(Avex-73) |  | 3.7 x 10<sup>-6</sup> | 2.4 | 49.5±7.9<sup>ab</sup> | 67.7±8.6<sup>ab</sup> |
| 22 | Benzoyl Peroxide |  | 4.2 x 10<sup>-4</sup> | 3.1 | 81.8±4.8<sup>a</sup> | Not tested |

\* For each dose, different letters indicate statistically significant differences between tested compounds (p < 0.05), Mann–Whitney U test with Bonferroni correction (adjusted p values a = 0.0071)
**NA:** Fluorescence Intensity decrease <50% under assay conditions (0-60 μM ligand)
**ND:** Interference with 1-NPN

To explore novel ligand scaffolds and lead structures retaining measurable repellent activity for further optimization, we assessed whether increasing the dose could enhance the activity of compounds exhibiting low-to-mild repellency. Accordingly, compounds #3, #5, #12, #14, #15, and #18 were subsequently tested at 0.4 mg/cm².

As shown in Table 4, increasing the dose did not significantly improve the repellent activity of o,p′-DDE (#14) or compounds #12 and #15, which remained in the low-to-mild activity range (statistical group b). In contrast, 2-phenylethyl benzoate (#5) and Avex-73 (#18) showed substantially increased repellency, reaching RIs close to or above 60%, thereby identifying these compounds as promising scaffolds for further optimization. Notably, fluorene (#3) exhibited a marked dose-dependent increase in activity, reaching an RI of 79.2% at 0.4 mg/cm², with a statistically significant difference (group a).

Overall, evaluation of the seven docking hits revealed several promising chemotypes. Benzoyl peroxide (#22), an FDA-approved active ingredient in over-the-counter topical acne products, emerged as the most potent repellent and one of the strongest AgamOBP4 binders. Avex-73 (#18), an advanced clinical-stage drug candidate, also combined strong AgamOBP4 binding with substantial repellent activity. In addition, two compounds that exhibited pronounced repellent activity at the higher dose retrieved from FooDB. Fluorene (#3) which its occurrence in food databases may reflect environmental or processing-related contamination, and 2-Phenylethyl benzoate (#5), (FEMA No. 2860), which is listed by the U.S. FDA under 21 CFR §172.515 as a synthetic flavoring substance/adjuvant.

### AI-based modelling of ligands binding

To gain insight into the molecular basis of the observed *in vitro* and *in vivo* activities, AI-based structural modeling was performed for all seven docking hits, enabling comparative analysis of their predicted binding modes within the AgamOBP4 cavity.

### Validation of the Protenix prediction protocol

The Protenix prediction protocol was validated by testing whether the protocol could reproduce the binding mode of the co-crystallized ligands 1-NPN and Indole. The predicted 1-NPN binding pose closely reproduced the crystallographic binding mode with a ligand heavy-atom RMSD of 1.109 Å. Similarly, the predicted IND1 pose showed near-complete overlap with the crystallographic pose (RMSD, 1.148 Å). In contrast, IND2 showed only partial overlap with its crystallographic pose (RMSD, 3.907 Å). Furthermore, the Protenix-predicted poses (**Figure 6**) reproduce most of the crystallographically observed protein-ligand contacts (**Figure 1B-I**, **Figure 2A-I** and **2A-II)**. These results show that the protocol successfully restored the experimentally characterized binding mode and can therefore be used to generate plausible poses for the seven compounds for which the exact crystallographic structures with AgamOBP4 have not yet been experimentally determined.

**Figure 6.**
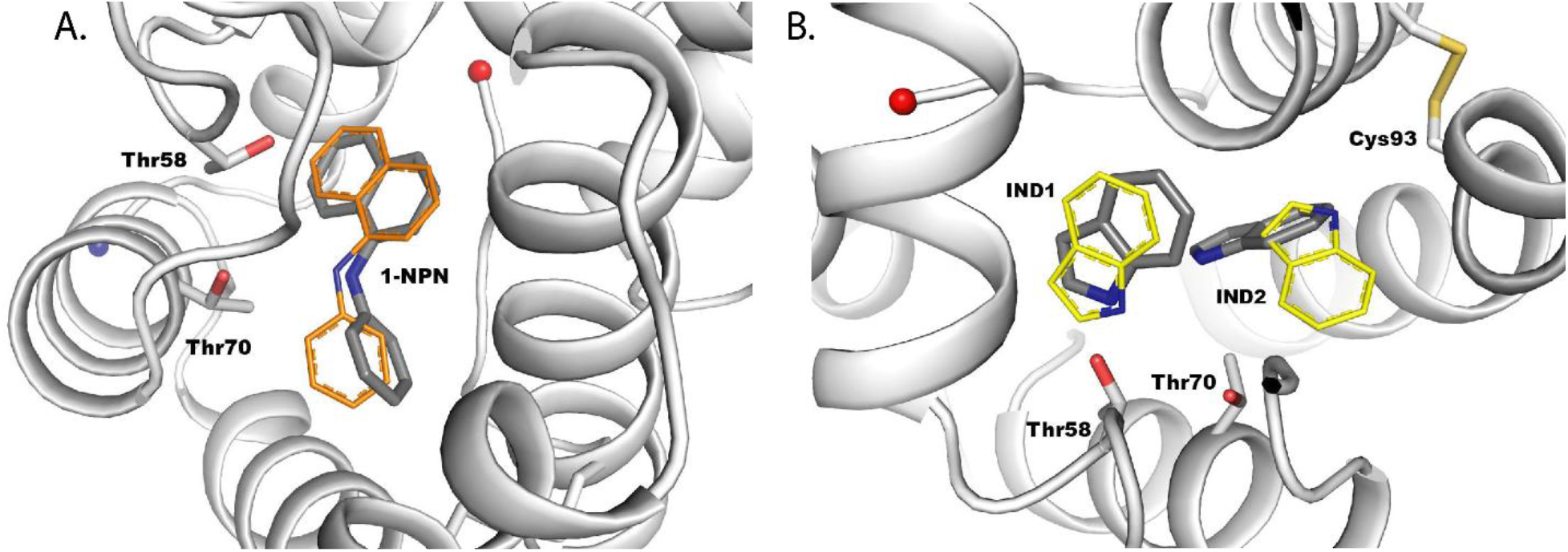
Superimposition of the Protenix-predicted (A) AgamOBP4–1–NPN and (B) AgamOBP4–2xIndole structures onto the corresponding crystallographic structures. The Protenix-predicted AgamOBP4 structure and ligand poses are shown in grey, while the crystallographic 1-NPN and Indole poses are shown in orange and yellow, respectively. The key interacting residues Thr70 and Thr58 are shown as sticks.

### Protenix Modeling of ligands’ binding modes

The predicted AgamOBP4–ligand complexes exhibited high confidence scores, including high pTM, ipTM, and pLDDT values, as well as low GPDE values (**Supplementary Table S1**). Nevertheless, the reliability of the predicted binding poses was assessed primarily by comparison with the experimentally determined ligand-binding site, binding mode, and crystallographic protein-ligand interactions, rather than by confidence scores alone.

The predicted binding pose of ligand **#3 (Fluorene)** showed only partial overlap with the crystallographic position of 1-NPN **(Figure 7A, left**), potentially accounting for its limited ability to displace 1-NPN in the fluorescence competitive binding assays (**Figure 5** and **Table 3**). In this predicted complex, the ligand forms 25 contacts, including π-alkyl interactions with residues Ala53, Leu90, and Ala111, as well as a π–π T-shaped interaction with the C-terminal residue Phe124 (**Figure 7A, right**).

**Figure 7.**
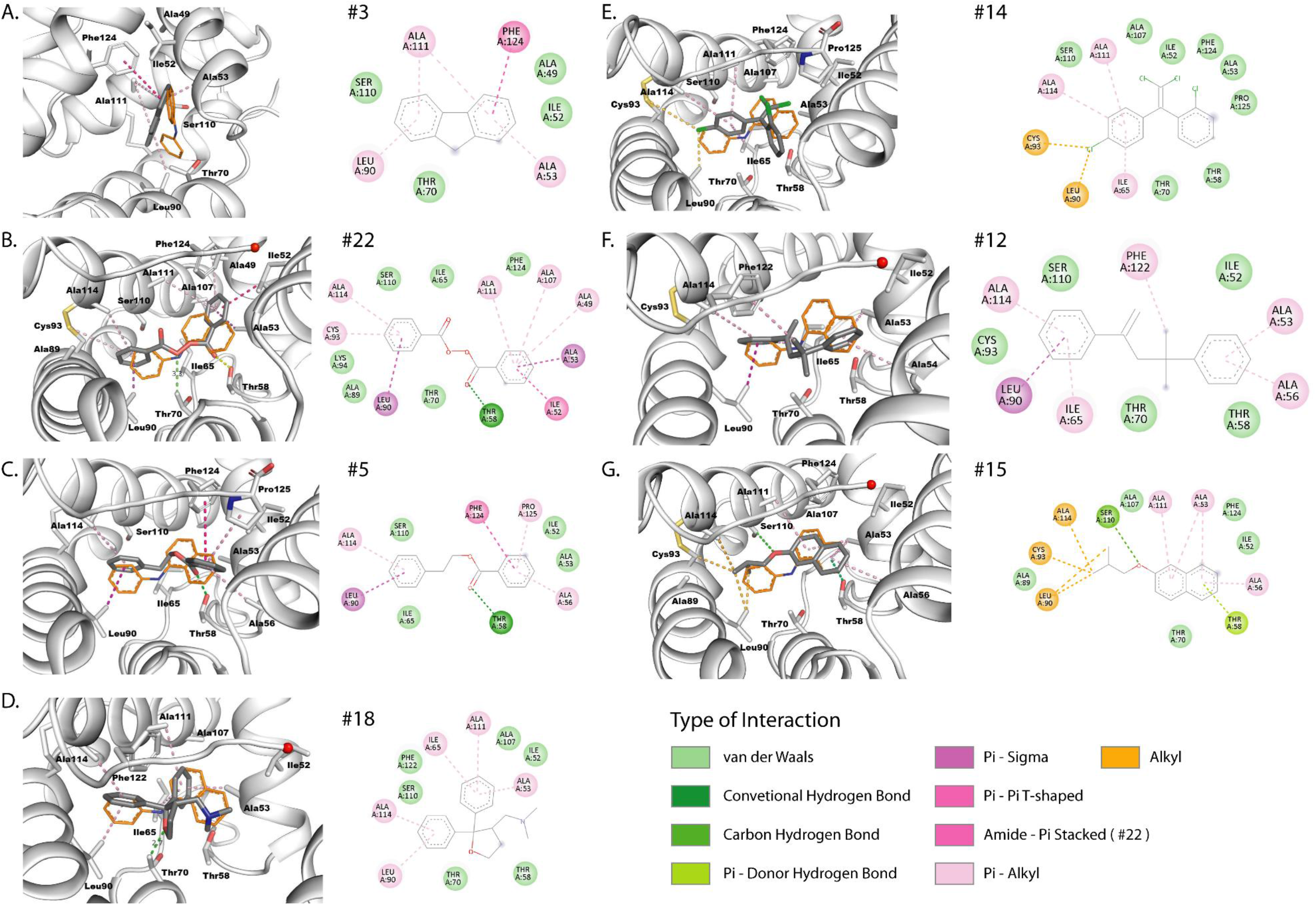
AI-based predicted binding modes. The structural panels depict superimposition of the Protenix-predicted AgamOBP4-ligand complexes (grey) onto the crystallographic AgamOBP4-1-NPN structure, with the crystallographic 1-NPN shown in orange. For each compound, the predicted binding mode within the AgamOBP4 binding cavity (left) and the corresponding 2D interaction diagram (right) are shown side by side. Binding-site residues located within 4 Å of the ligands are shown as sticks and spheres colored by interaction type in the corresponding 2D interaction diagrams.

In contrast, the predicted binding poses of all other ligands exhibited substantial overlap with the experimentally determined positions of 1-NPN and the two indole molecules in the crystal structure (**Figure 7B-7F**).

Among the investigated ligands, the diphenyl compound **#22 (benzoyl peroxide)** was predicted to adopt a binding pose in which one phenyl ring is oriented nearly coplanar with the naphthalene ring of 1-NPN and IND1 (22,9° and 16.7°, respectively), while its second phenyl ring is aligned with the phenyl group of 1-NPN and IND2, displaying interplanar angles of 43.5° and 18.7°, respectively. Consequently, benzoyl peroxide exploits its two aromatic groups to bind to the two subsites in a manner that closely resembles the experimentally determined ligand-binding modes. In its predicted pose, benzoyl peroxide makes 60 contacts, including a hydrogen bond (2.49 Å) between its O4 carbonyl oxygen and the hydroxyl group of Thr58. This interaction reproduces the π-donor hydrogen bond observed between Thr58 and the naphthalene ring of 1-NPN or the IND1 molecule in the crystal structures. It is worth noting that the O2 peroxide oxygen lies 3.3 Å from the CG2 atom of the Thr70 side chain (**Figure 7B, left)**. Although the current model does not predict a direct hydrogen bond to Thr70, rotation around the Cα– Cβ bond of the threonine side chain could bring its OG1 hydroxyl group into a favorable position for hydrogen bond formation with the ligand’s peroxide oxygen atom. Such an interaction, if established, would be analogous to the key hydrogen bond observed between Thr70 and either the amine nitrogen of 1-NPN or the IND1 nitrogen in the corresponding crystal structures. The high number of intermolecular contacts is further complemented by aromatic interactions involving the phenyl rings, including π-σ interactions with Ala53 and Leu90, π-alkyl interactions with Ala49, Ala107, Ala111, Ala114, and Cys93, and an amide-π stacking interaction with Ile52 (**Figure 7B, right**). In addition, several close vdW contacts were predicted between atoms C12 and C13 of one phenyl ring and the CE1 and CZ atoms of Phe124, with interatomic distances ranging from 2.65 to 3.67 Å.

#### Ligand #5 (2-phenylethyl benzoate)

similar to benzoyl peroxide (#22), contains two phenyl rings that occupy the experimentally defined aromatic subsites. However, unlike benzoyl peroxide, only one of its two aromatic moieties closely aligns with the aromatic systems of 1-NPN and the indole molecules (**Figure 7C, left)**. Specifically, its phenethyl ring is inclined by 45.0° and 23.3° relative to the planes of the 1-NPN phenyl group and IND2, respectively, closely resembling the orientation adopted by the corresponding phenyl ring of benzoyl peroxide in subsite 2. This positioning enables the phenethyl ring of ligand #5 to retain two aromatic contacts analogous to those observed for the benzoyl ring of ligand #22 (**Figure 7B, right**), namely a π–σ interaction with Leu90 and a π-alkyl interaction with Ala114 (**Figure 7C, right**). In contrast, the benzoate phenyl ring is positioned above and nearly perpendicular to the plane of the 1-NPN naphthalene moiety and IND1, forming interplanar angles of 79.0° and 84.8°, respectively. Consequently, unlike ligand #22, the benzoate moiety engages in π–π T-shaped and π-alkyl interactions with a different set of residues surrounding subsite 1, including Ala56 and the C-terminal aminoacids Phe124 and Pro125. Notably, the carbonyl oxygen of compound #5 participates in a hydrogen bond with Thr58, reproducing the key interaction observed for benzoyl peroxide (#22). However, due to its lower number of oxygen atoms available for hydrogen bonding, 2-phenylethyl benzoate lacks the potential to form a hydrogen bond with Thr70. Overall, despite possessing a comparable molecular length and aromatic framework, 2-phenylethyl benzoate is predicted to be less efficiently packed within the binding pocket, engage in more distant contacts with Phe124, and establish only 25 interactions, compared with the 60 of benzoyl peroxide. These differences, together with the higher flexibility conferred by the ethylene linker connecting the two aromatic rings of 2-phenylethyl benzoate, which allows greater conformational freedom, may account for the experimentally observed variation in the binding affinities of the two compounds (**Figure 5 and Table 3**).

#### Ligand #18 (Avex-73)

was predicted to adopt a binding pose in which one phenyl ring is oriented nearly perpendicular to the naphthalene moiety of 1-NPN and IND1, whereas its second phenyl ring is tilted by 69.5° and 44.0° relative to the phenyl ring of 1-NPN and IND2, respectively (**Figure 7D, left**). In this orientation, the ligand participates in 24 contacts, including π-alkyl interactions of its phenyl rings with Ala53, Ile65, Leu90, Ala111, and Ala114. (**Figure 7D, right**). It is worth noting that the oxygen atom of the tetrahydrofuran ring of ligand #18 lies 2.71 Å from the CG2 atom of Thr70, potentially allowing the formation of a hydrogen bond with the Thr70 hydroxyl group upon rotation of its side chain about the Cα-Cβ bond (**Figure 7D, left**).

#### Ligand #14 (o,p′-DDE)

is predicted to form 25 interactions. It contains a gem-dichlorovinyl group (–CCl₂=CH–) that is positioned above, yet in close proximity to the 1-NPN naphthalene moiety and IND1, exhibiting interplanar angles of 20.3° and 36.9°, respectively. However, the ortho-chlorophenyl group lies outside the IND1 plane and engages only in vdW interactions with residues in subsite-1. In contrast, the para-chlorophenyl ring occupies aromatic subsite-2 and closely aligns with the phenyl ring of 1-NPN and IND2, with interplanar angles of 25.2° and 5.8°, respectively (**Figure 7E, left**). In this position, the p-ring participates in π-alkyl interactions with Ala65, Ala111, and Ala114, while one of its chlorine atoms forms two alkyl contacts with Leu90 and Cys93, thereby resembling the aromatic interaction pattern observed in the IND2 molecule (**Figure 7E, right**).

Among the tested compounds containing two distinct phenyl rings, compound **#12 (4-methylpent-1-ene-2,4-diyl dibenzene)** is the only one that lacks functional groups capable of mediating direct hydrogen-bonding or other specific polar interactions with the binding pocket. In total, compound #12 is predicted to establish 28 contacts. Its alkene-associated phenyl ring is modeled to overlap almost perfectly with the phenethyl phenyl ring of ligand #5 (2-phenylethyl benzoate) within subsite 2. On the other hand, its tertiary alkyl phenyl ring occupies subsite 1, adopting an orientation nearly perpendicular to the aromatic planes of 1-NPN and IND1 (85.5° and 86.3°, respectively) (**Figure 7F, left**), resembling the predicted arrangement of the benzoate phenyl ring of compound #5 (**Figure 7C, left**). However, their linkers follow distinct trajectories through the binding pocket. Consequently, the tertiary alkyl phenyl ring of compound #12 and the benzoate phenyl ring of compound #5 are rotated by approximately 45° relative to each other, resulting in different patterns of aromatic interactions within subsite 1. Specifically, the modeled pose suggests that compound #12 retains the π-alkyl interaction with Ala56, but lacks π-type contacts with the C-terminal residues Phe124 and Pro125. Instead, its branched alkyl linker forms a π-alkyl interaction with Phe122 (**Figure 7F, right**).

Finally, compound **#15 (2-isobutoxynaphthalene)** is unique among the selected hits as it contains only a single aromatic naphthalene system linked to an isobutoxy substituent. As a result, it lacks the second aromatic moiety, characteristic of the other compounds and enables the simultaneous occupation of both aromatic subsites. The compound was predicted to adopt a binding mode in which its naphthalene group closely overlaps with the naphthalene ring of 1-NPN and IND1 in the aromatic subsite 1. The isobutoxy substituent extends toward subsite 2, where one CH-CH₃ fragment of its isopropyl tail lies in a position analogous to that occupied by the two adjacent aromatic carbon atoms of the 1-NPN phenyl ring or the pyrrolic N1-C8 edge of IND2 (**Figure 7G, left**). Notably, despite lacking a second aromatic ring, compound #15 is predicted to establish 34 contacts, exceeding the number of interactions for the other examined compounds (24-28 contacts), except for benzoyl peroxide (#22), which is predicted to participate in 60 contacts. This can be attributed to the extended interaction network developed by its isobutoxy substituent in subsite 2, including a vdW contact with Ala89 and alkyl interactions with Leu90, Cys93, and Ala114 (**Figure 7G, right**). In addition, the ether oxygen is predicted to engage in a carbon-oxygen interaction with the Cβ atom of Ser110 (3.66 Å), classified by Discovery Studio 2025 analysis as a C-H···O hydrogen bond. Although these interactions are weaker than conventional hydrogen bonds, they have been proposed to contribute to the stabilization of protein-ligand complexes, especially when stronger hydrogen bonds are absent (Baures, Wiznycia et al., 2000, Musah, Jensen et al., 1997, Sarkhel & Desiraju, 2004). Notably, the two alternative rotamers of the modeled Ser110 side chain, generated by rotation about the Cα-Cβ bond as implemented in Coot, position the OG hydroxyl group within hydrogen-bond distances of 2.88 Å and 3.16 Å to the ether oxygen. In subsite 1, the pattern of aromatic interactions mediated by the naphthalene ring closely resembles the one formed by the crystallographic IND1 (**Figure 2A-I**). This includes π-alkyl interactions with Ala53, Ala56, and Ala111, as well as a π-donor hydrogen bond with the key residue Thr58, also observed in both the 1-NPN and indole crystal structures (**Figure 1B-I and Figure 2A-I)**.

## Discussion

The fluorescent probe 1-NPN, found bound to AgamOBP4, exhibits a compact architecture in which the phenyl and naphthalene rings are connected via a single amine linkage, resulting in restricted conformational freedom and a well-defined geometry that is well-suited for its efficient binding to the AgamOBP4 cavity. Crystallographic analysis revealed that the AgamOBP4 cavity is spacious and largely preformed, allowing accommodation of either a single extended ligand or multiple aromatic molecules simultaneously. While 1-NPN occupies the bottom of the cavity as a single molecule, the indole crystal structure revealed two molecules bound at neighboring subsites. Superposition of the complexes showed that the two indoles closely overlap with the naphthalene and phenyl moieties of 1-NPN, indicating that the cavity contains two distinct aromatic-recognition regions. The primary binding subsite (IND1 and 1-NPN naphthalene ring subsite) appears to be the preferred indole-binding region, as the AgamOBP4 structure 3Q8I revealed indole occupancy only at this site. In contrast, in this study ligand binding at the secondary binding subsite (IND2 and 1-NPN phenyl ring subsite) was observed only after soaking the protein crystals with 25 mM indole, suggesting that binding to this subsite likely requires substantially higher ligand concentrations.

Both crystal structures identified Thr70 and Thr58 as key residues involved in ligand recognition, mediating conventional hydrogen bonds and π-donor hydrogen-bond interactions. Notably, ligand binding is accompanied by only limited conformational rearrangements, indicating that AgamOBP4 recognizes odorants through a largely preformed binding cavity rather than by substantial induced-fit adaptations, consistent with observations reported for several other OBP-ligand complexes (Christodoulou et al., 2026, Drakou et al., 2017, Liggri et al., 2023b, Tsitsanou, Thireou et al., 2012). Together, these findings reveal that AgamOBP4 possesses a dual aromatic binding architecture capable of recognizing structurally diverse aromatic ligands and further provide a structural framework for the discovery of novel host-seeking disruptors targeting this protein.

In contrast to the local structural conservation of the AgamOBP4 ligand-binding pocket, pH-dependent conformational changes involving the flexible N-terminal region (residues 1-9) and the C-terminal tail (residues 120-125), including pronounced differences in the relative orientations of Phe122 and Phe124, have been previously documented (Mam et al., 2023). To account for this conformational variability, the docking campaign was performed against all available AgamOBP4-ligand structures, including the indole complex determined at pH 6.85 (PDBid: 3Q8I) and the two structures determined at pH 4.6 (this study, PDBids: 34XM and 34XN). Interestingly, the 1-NPN-bound structure at pH 4.6 emerged as the preferred docking template for all selected hits, with the sole exception of compound #14, which exhibited a more favorable docking score against the 3Q8I structure (**Table 2**). AgamOBP4 is exposed to a complex pH landscape extending from the negatively charged sensillum pore wall, through the near-neutral sensillum lymph, toward the more acidic environment surrounding the olfactory receptors. Consequently, although the identified ligands were discovered using a low-pH structural model, the binding affinities reported in this study were determined at pH 8.0, and the crystallographic positions of IND1 at pH 4.6 and pH 6.85 closely overlap, indicating that their interaction with AgamOBP4 is unlikely to be restricted to a single conformational state. This hypothesis is consistent with the overall conservation of the ligand-binding cavity across the available AgamOBP4 structures. Accordingly, these compounds should be viewed as AgamOBP4-targeting ligands capable of interacting with the protein across different stages of OBP function, including capture near the sensillum pore, transport through the sensillum lymph, and release near olfactory receptors. Additionally, they may facilitate protein-protein interactions.

Furthermore, because mosquito olfaction relies on the detection of airborne semiochemicals, the incorporation of a volatility filter was an essential element of the screening workflow. This step ensured the selection of hits possessing sufficient vapor pressure to be detected in the air and to participate in olfaction-mediated signaling, while excluding excessively volatile compounds that could compromise duration and efficacy during application (Gupta & Bhattacharjee, 2007, Liggri et al., 2023a).

The structure-based virtual screening pipeline employed in this work proved highly effective in identifying strong AgamOBP4 binders, as demonstrated by the subsequent *in vitro* validation of selected candidate ligands.

Among the compounds experimentally evaluated, o,p′-DDE (#14), Avex-73 (#18), and benzoyl peroxide (#22) emerged as the strongest AgamOBP4 binders (**Figure 5 and Table 3**). Despite their distinct chemical scaffolds, all three compounds share a common architectural feature: two appropriately spaced aromatic moieties connected by rigid or semi-rigid linkers. Molecular modeling indicated that these preorganized scaffolds are well suited to simultaneously occupy the two experimentally defined aromatic subsites, thereby reproducing key structural features of the binding modes observed for 1-NPN and the indole molecules.

In contrast, compound #5 (2-phenylethyl benzoate), which contains an extended O-CH₂-CH₂-ester linker, can adopt a wider range of conformations in solution. This increased conformational flexibility is expected to incur a larger entropic penalty upon binding and may hinder the optimal simultaneous fit of its aromatic rings within the two aromatic subsites, thereby leading to a lower binding affinity for AgamOBP4 (**Figure 5 and Table 3**). Nevertheless, its moderate affinity is comparable to that previously reported for indole (15 to 26 μΜ) (Mam et al., 2023, Qiao et al., 2010), warranting its further investigation in behavioral assays.

As discussed extensively elsewhere (Christodoulou et al., 2026, Murphy, Booth et al., 2013), fluorescence probe-displacement assays provide only an indirect measure of ligand binding and are subject to several inherent limitations. The analysis assumes that the tested ligand and the fluorescent probe occupy the same binding site and that binding follows a 1:1 equilibrium model. Deviations from these assumptions may affect the accuracy of the calculated IC₅₀ and Kᵢ values, underscoring the importance of integrating fluorescence binding data with structural analyses to obtain a more comprehensive assessment of ligand recognition. Furthermore, reliable affinity determination requires that the tested ligand neither interferes with probe fluorescence nor exhibits significant intrinsic fluorescence under the assay conditions (Feng, Fei et al., 2025, Leal & Leal, 2015). Therefore, docking hits such as fluorene (#3), which showed only marginal displacement of 1-NPN (**Figure 5**), likely due to its partial overlap with the crystallographic position of 1-NPN (**Figure 7A, left**), should not be excluded from further behavioral evaluation. The same applies to 4-methylpent-1-ene-2,4-diyl dibenzene (#12) and 2-isobutoxynaphthalene (#15), both of which interfered with the fluorescence assay (**Figure 4**).

Although high affinity for an OBP is desirable for detecting low concentrations of semiochemicals in the headspace, binding affinity alone may not fully determine biological activity. OBPs are not merely nonspecific solubilizers of hydrophobic molecules but function as physiological transport proteins within the olfactory system, contributing to its remarkable selectivity and sensitivity through their coordinated interplay with odorant receptors (ORs) (Forstner, Breer et al., 2009, Leal, 2005). It has therefore been proposed that, for efficient receptor activation, a ligand must not only bind an OBP but also be presented to the OR in a physicochemically compatible conformation. Consequently, OBP and OR binding pockets are expected to share, at least in part, common structural and chemical features, including complementary size, shape, hydrophobic properties, and spatial arrangements of key interaction determinants. Thus, ligand binding mode was considered an important selection criterion alongside volatility, docking score, and predicted binding affinity, as biologically active molecules are expected not only to bind OBPs efficiently but also to adopt OBP-bound conformations that preserve the molecular features required for their subsequent recognition by the ORs (Drakou et al., 2017, Kritsi, Liggri et al., 2022, Liggri et al., 2023a, Murphy et al., 2013, Thireou et al., 2018, Tsitsanou et al., 2012, Zographos, Eliopoulos et al., 2018).

Analysis of ligand-binding modes predicted by AI-based modeling suggested that simultaneous occupation of both aromatic subsites by aromatic moieties mimicking the orientations of IND1 and IND2 molecules, and the formation of conventional and π-donor hydrogen bonds with residues Thr70 and Thr58, may be associated with high repellent activity.

Consistent with this hypothesis, benzoyl peroxide (#22), which closely reproduced this binding pattern, displayed strong repellency (RI = 81.8% at 0.2 mg/cm²). In contrast, 2-phenylethyl benzoate (#5) and Avex-73 (#18) adopted markedly different binding modes within the primary aromatic subsite 1. In both compounds, the aromatic group was positioned approximately perpendicular to the aromatic plane of IND1. In addition, 2-phenylethyl benzoate (#5) lacked the hydrogen bond with Thr70, whereas Avex-73 (#18) failed to establish the π-donor hydrogen bond with Thr58, both characteristic interactions of IND1. The absence of these key binding-mode features may contribute to their lower repellent activity relative to benzoyl peroxide (#22). Behavioral assay results further supported this interpretation, as neither compound exceeded the threshold for significant repellency (arbitrarily defined as RI ≥ 60%) at the standard dose (0.2 mg/cm²). Both ligands exhibited only moderate activity at 0.4 mg/cm², with RI values of 57.5% and 67.7% for 2-phenylethyl benzoate and Avex-73, respectively. It should also be noted that the relatively low volatility of Avex-73, evidenced by a vapor pressure of 3.7 x 10⁻⁶ mmHg, may have further contributed to its reduced repellency at the lower dose (0.2 mg/cm²) by limiting the availability of airborne molecules.

Among the remaining ligands, compound 4-methylpent-1-ene-2,4-diyl dibenzene (#12), which shares the perpendicular aromatic-ring orientation observed for the moderately active 2-phenylethyl benzoate (#5) and Avex-73 (#18), and o,p′-DDE (#14), whose aromatic groups closely mimic the arrangement displayed by the highly active benzoyl peroxide (#22), share a common characteristic: neither is capable of establishing the key hydrogen-bond interactions with Thr70 and Thr58. In contrast, compound 2-isobutoxynaphthalene (#15) largely preserves the orientation of the IND1 within the primary aromatic subsite and retains the π-donor hydrogen bond with Thr58, but lacks a second aromatic moiety capable of occupying the secondary aromatic subsite. None of these compounds exhibited significant repellency, even at the highest tested dose (0.4 mg/cm²).

Collectively, these findings reinforce the hypothesis that potent repellency requires a combination of dual-subsite occupancy, appropriate aromatic-ring orientation, and the formation of key interactions with Thr70 and Thr58, rather than any of these features alone. Moreover, these results show that favorable binding affinity alone is insufficient to predict behavioral activity. For example, compounds such as o,p′-DDE (#14) and Avex-73 (#18) exhibited high binding affinities but only limited or moderate repellent activity at the higher dose tested. Taken together, these observations suggest that biological activity depends not only on the strength of ligand binding but also on the specific binding mode adopted within the AgamOBP4 cavity.

Interestingly, fluorene (#3), which does not appear to satisfy any of the proposed binding-mode criteria in its top-ranked pose, exhibited strong repellent activity at the highest tested dose (RI = 79.2% at 0.4 mg/cm²). This apparent discrepancy suggests that the biologically relevant binding mode may differ from the top-ranked prediction. Indeed, inspection of the four lower-ranked poses among the five generated models (**Supplementary Table S2**), revealed alternative conformations that more closely resemble the spatial arrangement of the crystallographic ligands. In these alternative poses, the fluorene scaffold is positioned to establish a π-donor hydrogen bond with the key residue Thr58. Furthermore, alternative Thr70 rotamers position the OG1 hydroxyl group within π-donor hydrogen-bonding distance of the aromatic system located at the secondary subsite 2 (**Supplementary Figure S4**). Therefore, it cannot be excluded that, under physiological conditions, fluorene may adopt an alternative orientation that reproduces one or more of the key structural and interaction features identified herein, including occupation of the aromatic subsites and interactions with Thr70 and/or Thr58. This possibility remains to be investigated in future crystallographic studies and may provide a structural rationale for the unexpected, repellent activity of fluorene.

Overall, the results demonstrate that the discovery pipeline employed herein represents an efficient and cost-effective approach for the identification of novel high-affinity AgamOBP4 ligands. Moreover, the use of screening libraries comprising approved drugs and food-derived compounds enabled the discovery of structurally diverse ligand scaffolds recognized by AgamOBP4, highlighting the potential of drug-repurposing strategies for the identification of behaviorally active molecules targeting the insect olfactory system. Such compounds may benefit from existing safety, toxicological, and regulatory data, potentially facilitating their further development into safe products. The availability of pre-existing safety and regulatory information may reduce the extent of additional testing required, thereby potentially decreasing the time and cost associated with product development compared to entirely novel chemical entities, whether synthetic or naturally derived. This consideration is particularly relevant in the biocides sector, where regulatory approval under frameworks such as the U.S. Federal Insecticide, Fungicide, and Rodenticide Act (FIFRA) (https://www.epa.gov/pesticides) and the EU Biocidal Products Regulation (BPR) (ECHA, 2024) may necessitate substantial financial investment in testing and several years of regulatory review. For example, benzoyl peroxide (#22), identified in this study as one of the strongest AgamOBP4 binders and exhibiting significant repellent activity, is an FDA-approved active ingredient widely used in topical acne treatments. Its extensive existing safety, toxicological, formulation, and manufacturing data, make it an attractive candidate for further development as a mosquito repellent for topical human application.

Overall, these findings demonstrate that our structure-guided screening strategy can identify behaviorally active compounds across the diverse chemical space, including established pharmaceutical ingredients and compounds used in flavor and fragrance applications, providing promising starting points for the development of novel mosquito repellents.

Furthermore, recognition of environmental pollutants such as o,p′-DDE and fluorene opens new opportunities for the development of AgamOBP4-based biosensors for food-quality and environmental monitoring applications. Moreover, identifying o,p’-DDE, a persistent metabolite of the insecticide DDT, as one of the strongest AgamOBP4 ligands may have implications beyond olfaction. Growing evidence indicates that several insect chemosensory proteins (CSPs), and to a lesser extent OBPs, can interact with insecticides and are involved in sequestration, detoxification-associated processes, and insecticide resistance mechanisms (Tsouri & Douris, 2025). Therefore, this finding suggests that AgamOBP4 may participate in recognizing insecticides in addition to its established role in odorant perception.

This methodology could be readily adapted and extended to other mosquito molecular chemosensory targets for which structural information, such as crystal or cryo-EM structures, is available, including OBPs, CSPs, and ORs, implicated in host-seeking behavior. Such an approach could accelerate the discovery of next-generation repellents/attractants for the prevention and control of vector-borne diseases.

## Acknowledgments

This research was supported by funding from the project “iStink – Insect-Inspired Biosensor for Water Quality Monitoring” under the “Exploratory Interdisciplinary Projects of Early-Career Researchers, 1st NHRF Internal Call 2024-2026” (Project Number: NHRF-0867-4). We also acknowledge the support provided to the X-ray protein crystallography facility at the National Hellenic Research Foundation (NHRF) by the project “INSPIRED – The National Research Infrastructures on Integrated Structural Biology, Drug Screening Efforts and Drug Target Functional Characterization” (MIS 5002550). Access to the EMBL Hamburg DESY/DORIS III Synchrotron Radiation Facility, Germany, was supported by FP7 project “BioStruct-X” (GA-283570). We extend our gratitude to the staff of beamline X13 for their assistance. The authors thankfully acknowledge RES resources provided by Barcelona Supercomputing Center in MareNostrum5 to BCV-2025-3-0024. Powered@NLHPC. This research was also partially supported by the super-computing infrastructure of NLHPC (ECM-02). We also thank Stratis Zografos (School of Architecture, Technical University of Crete, Greece) for his assistance in the design and preparation of the graphical artwork presented in Figure 3 and the Graphical Abstract.

## Supporting Information description

- Supplementary Figures and Tables: **Supporting Information.docx**

## Ethics statement

The laboratory strain of *Ae. albopictus* used in this study was established using mosquito eggs collected from ovitraps from different areas in Greece. The collection areas were public and not privately owned or protected. Mosquito egg collections from the field did not involve endangered or protected animal species. Consequently, the establishment of the laboratory mosquito strain did not require a specific permit. The repellence studies abide by the Declaration of Helsinki principles. The Ethics Committee of Benaki Phytopathological Institute concluded that the current study was implemented in accordance with the Ethics Code for Research.

## Disclosure of AI usage

During the preparation of this work, the authors used the AI-based deep-learning model **protenix_base_constraint_v0.5.0**, available through the **Tamarind Bio platform** to generate models of the Protein-ligand complexes, and **Microsoft Copilot** and **Paperpal** for grammar checking and language editing. After using this tool/service, the authors reviewed and edited the content as needed and take full responsibility for the content of the published work.

## CRediT Authorship Contributions

**S.E.Z., H.P.-S., and K.E.T.**: Conceptualization.

**S.E.Z. and K.E.T.**: Funding acquisition, Project administration.

**S.E.Z. and K.E.T.**: Methodology (*in vitro* studies), Supervision.

**H.P.-S.**: Methodology (*in silico* studies), Supervision.

**E.C.V.S.**: Investigation (*in vitro* binding assays, AI-based modeling), Data curation, Formal analysis.

**J.-R.A.-F.**: Investigation (virtual screening), Data curation, Formal analysis.

**C.E.D., K.E.T., and S.E.Z.**: Investigation (crystallographic studies), Data curation, Formal analysis.

**A.M. and D.P.P.**: Methodology (*in vivo* behavioral assays), Supervision.

**E.C.V.S., M.M., and V.K.**: Investigation (*in vivo* behavioral assays), Data curation.

**D.P.P.**: Formal analysis (behavioral assay data).

**E.C.V.S. and S.E.Z.**: Formal analysis and interpretation of AI-based modeling results.

**E.C.V.S., J.-R.A.-F., and K.E.T.**: Writing - first draft.

**E.C.V.S., J.-R.A.-F., K.E.T. and S.E.Z**: Writing - original draft.

**All authors**: Writing - review & editing.

**All authors** have read and approved the final version of the manuscript.

## Abbreviations

OBP: Odorant binding protein
AgamOBP4: Odorant binding protein 4 from *Anopheles gambiae*
OR: odorant receptor
1-NPN: *N*-phenyl-1-naphthylamine
IND: Indole
IPTG: isopropyl-galacto-pyranoside
vdW: van der Waals force
RMSD: root mean square deviation

---

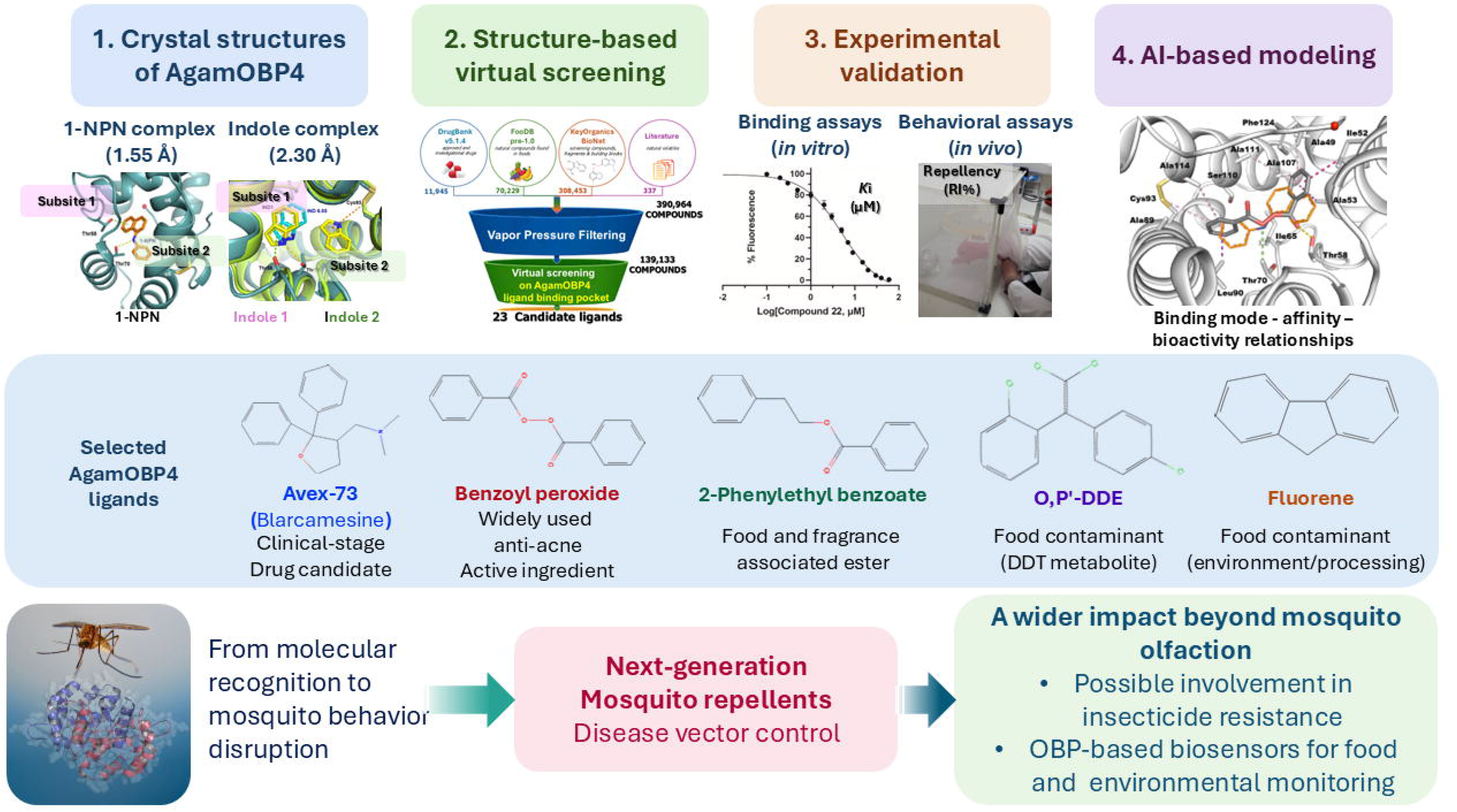

## References

Abendroth JA, Moural TW, Wei H, Zhu F (2023) Roles of insect odorant binding proteins in communication and xenobiotic adaptation. 3

Abramson J, Adler J, Dunger J, Evans R, Green T, Pritzel A, Ronneberger O, Willmore L, Ballard AJ, Bambrick J, Bodenstein SW, Evans DA, Hung CC, O’Neill M, Reiman D, Tunyasuvunakool K, Wu Z, Zemgulyte A, Arvaniti E, Beattie C et al. (2024) Accurate structure prediction of biomolecular interactions with AlphaFold 3. Nature 630: 493–500

Andronopoulou E, Labropoulou V, Douris V, Woods DF, Biessmann H, Iatrou K (2006) Specific interactions among odorant-binding proteins of the African malaria vector Anopheles gambiae. Insect Mol Biol 15: 797–811

Baures P, Wiznycia A, Beatty AJB, chemistry m (2000) Hydrogen bonding isosteres: bimolecular carboxylic acid and amine-N-oxide interactions mediated via CH··· O hydrogen bonds. 8: 1599–1605

Biessmann H, Andronopoulou E, Biessmann MR, Douris V, Dimitratos SD, Eliopoulos E, Guerin PM, Iatrou K, Justice RW, Kröber T, Marinotti O, Tsitoura P, Woods DF, Walter MF (2010) The Anopheles gambiae odorant binding protein 1 (AgamOBP1) mediates indole recognition in the antennae of female mosquitoes. PloS one 5: e9471

Biessmann H, Walter MF, Dimitratos S, Woods D (2002) Isolation of cDNA clones encoding putative odourant binding proteins from the antennae of the malaria-transmitting mosquito, Anopheles gambiae. Insect molecular biology 11: 123–32

Briand L, Nespoulous C, Huet JC, Takahashi M, Pernollet JC (2001) Ligand binding and physico-chemical properties of ASP2, a recombinant odorant-binding protein from honeybee (Apis mellifera L.). European journal of biochemistry 268: 752–60

Brito NF, Moreira MF, Melo ACA (2016) A look inside odorant-binding proteins in insect chemoreception. Journal of Insect Physiology 95: 51–65

ByteDance AMLAIST, Chen X, Zhang Y, Lu C, Ma W, Guan J, Gong C, Yang J, Zhang H, Zhang K, Wu S, Zhou K, Yang Y, Liu Z, Wang L, Shi B, Shi S, Xiao W (2025) Protenix - Advancing Structure Prediction Through a Comprehensive AlphaFold3 Reproduction. In bioRxiv

Chazapi E, Kritsi E, Potamitis C, Liggri PGV, Tsitsanou KE, Drakou CE, Michaelakis A, Papachristos DP, Zographos SE, Zervou M, Calogeropoulou T (2025) AgamOBP1-Directed Discovery of Repellents to Control the Spread of Mosquito-Borne Diseases. Chemmedchem

Christodoulou E, Stamati ECV, Saitta F, Papakyriakou A, Fessas D, Tsitsanou KE, Zographos SE (2026) Structural insights into ligand recognition by the pleiotropic odorant-binding protein AgamOBP9. International Journal of Biological Macromolecules 372: 153025

COLLABORATIVE COMPUTATIONAL PROJECT N (1994) The CCP4 suite: programs for protein crystallography. Acta Crystallogr D Biol Crystallogr 50: 760–3

Damberger FF, Michel E, Ishida Y, Leal WS, Wuthrich K (2013) Pheromone discrimination by a pH-tuned polymorphism of the Bombyx mori pheromone-binding protein. Proceedings of the National Academy of Sciences of the United States of America 110: 18680–18685

Davrazou F, Dong E, Murphy EJ, Johnson HT, Jones DN (2011) New insights into the mechanism of odorant detection by the malaria-transmitting mosquito Anopheles gambiae. J Biol Chem 286: 34175–83

Deng M, Xu X, Huang X, Xiao T, Wang W, Li J, Zhao X, Pan B, Jiang Y, He Z, Yang Z, Lu K (2025) Mechanistic exploration of odorant binding protein-mediated chlorpyrifos resistance in Nilaparvata lugens: Insights from insecticide sequestration and transcriptional regulation. International Journal of Biological Macromolecules 284: 138108

Drakou CE, Tsitsanou KE, Potamitis C, Fessas D, Zervou M, Zographos SE (2017) The crystal structure of the AgamOBP1*Icaridin complex reveals alternative binding modes and stereo-selective repellent recognition. Cell Mol Life Sci 74: 319–338

ECHA (2024) European Chemicals Agency, Practical guide on Union authorisation of biocidal products – February 2024. European Chemicals Agency,

Emsley P, Cowtan K (2004) Coot: model-building tools for molecular graphics. Acta Crystallographica Section D-Biological Crystallography 60: 2126–2132

Fan J, Francis F, Liu Y, Chen JL, Cheng DF (2011) An overview of odorant-binding protein functions in insect peripheral olfactory reception. Genetics and molecular research : GMR 10: 3056–69

FAO/IAEA Guidelines for routine colony maintenance of Aedes mosquito species v1.0. Vienna, Austria, 2017. In https://www.iaea.org/sites/default/files/21/06/nafa-ipc-manual-guidelines-for-routine-colony-maintenance-of-aedes-mosquito-species-v1.0.pdf

Feng M, Fei SG, Papakyriakou A, Christodoulou E, Lai WX, Luo WJ, Xia JM, Huang YG, Zographos SE, Swevers L, Sun JC (2025) Chemosensory protein 3 is a brain host factor for the induction of enhanced-locomotory activity in the BmNPV-silkworm infection model. Plos Pathogens 21

Forstner M, Breer H, Krieger J (2009) A receptor and binding protein interplay in the detection of a distinct pheromone component in the silkmoth Antheraea polyphemus. International journal of biological sciences 5: 745–57

French S, Wilson K (1978) Treatment of Negative Intensity Observations. Acta Crystallographica Section A 34: 517–525

Giatropoulos A, Papachristos DP, Kimbaris A, Koliopoulos G, Polissiou MG, Emmanouel N, Michaelakis A (2012) Evaluation of bioefficacy of three Citrus essential oils against the dengue vector Aedes albopictus (Diptera: Culicidae) in correlation to their components enantiomeric distribution. Parasitol Res 111: 2253–63

Gonzalez D, Rihani K, Neiers F, Poirier N, Fraichard S, Gotthard G, Chertemps T, Maïbèche M, Ferveur JF, Briand L (2020) The Drosophila odorant-binding protein 28a is involved in the detection of the floral odour ß-ionone. Cellular and molecular life sciences : CMLS 77: 2565–2577

Gupta RK, Bhattacharjee AK (2007) Discovery and Design of New Arthropod/Insect Repellents by Computer-Aided Molecular Modeling. In Insect repellents: Principles, methods, and uses, Debboun M FS, Strickman D (ed) pp 275–304. CRC, Boca Raton, FL

Halgren TA (1996) Merck molecular force field. 1. Basis, form, scope, parameterization, and performance of MMFF94. Journal of Computational Chemistry 17: 490–519

Han L, Zhang YJ, Zhang L, Cui X, Yu J, Zhang Z, Liu MS (2014) Operating mechanism and molecular dynamics of pheromone-binding protein ASP1 as influenced by pH. PloS one 9: e110565

Humphrey W, Dalke A, Schulten K (1996) VMD: Visual molecular dynamics. Journal of Molecular Graphics & Modelling 14: 33–38

Keil TA (1984) Surface Coats of Pore Tubules and Olfactory Sensory Dendrites of a Silkmoth Revealed by Cationic Markers. Tissue & Cell 16: 705–717

Kritsi E, Liggri PGV, Stamati ECV, Tsitsanou KE, Zographos SE, Michaelakis A, Papachristos D, Zoumpoulakis P (2022) A Combined Computational Methodology for the Discovery of Hit Compounds with Putative Insect Repellency Properties. ChemMedChem

Kruskal WH, Wallis WA (1952) Use of Ranks in One-Criterion Variance Analysis. J Am Stat Assoc 47: 583–621

Kythreoti G, Thireou T, Karoussiotis C, Georgoussi Z, Liggri PGV, Papachristos DP, Michaelakis A, Karras V, Zographos SE, Schulz S, Iatrou K (2024) Natural volatiles preventing mosquito biting: An integrated screening platform for accelerated discovery of ORco antagonists. J Biol Chem 300: 107939

Lagarde A, Spinelli S, Tegoni M, He X, Field L, Zhou JJ, Cambillau C (2011) The crystal structure of odorant binding protein 7 from Anopheles gambiae exhibits an outstanding adaptability of its binding site. J Mol Biol 414: 401–12

Leal GM, Leal WS (2015) Binding of a fluorescence reporter and a ligand to an odorant-binding protein of the yellow fever mosquito, Aedes aegypti. F1000Res 3: 305

Leal WS (2003) 15 - Proteins that make sense. In Insect Pheromone Biochemistry and Molecular Biology, Blomquist G, Vogt R (eds) pp 447–476. San Diego: Academic Press

Leal WS (2005) Pheromone reception. Chemistry of Pheromones and Other Semiochemicals Ii 240: 1–36

Leal WS, Ishida Y, Pelletier J, Xu W, Rayo J, Xu X, Ames JB (2009) Olfactory Proteins Mediating Chemical Communication in the Navel Orangeworm Moth, Amyelois transitella. PloS one 4: e7235

Leslie AGW (1992) Recent changes to the MOSFLM package for processing film and image plate data. Jnt CCP4/ESF-EACBM Newsl Protein Crystallogr No. 26

Liggri PGV, Perez-Garrido A, Tsitsanou KE, Dileep KV, Michaelakis A, Papachristos DP, Perez-Sanchez H, Zographos SE (2023a) 2D finger-printing and molecular docking studies identified potent mosquito repellents targeting odorant binding protein 1. Insect Biochem Mol Biol 157: 103961

Liggri PGV, Tsitsanou KE, Stamati ECV, Saitta F, Drakou CE, Leonidas DD, Fessas D, Zographos SE (2023b) The structure of AgamOBP5 in complex with the natural insect repellents Carvacrol and Thymol: Crystallographic, fluorescence and thermodynamic binding studies. Int J Biol Macromol 237: 124009

Mam B, Tsitsanou KE, Liggri PGV, Saitta F, Stamati ECV, Mahita J, Leonis G, Drakou CE, Papadopoulos M, Arnaud P, Offmann B, Fessas D, Sowdhamini R, Zographos SE (2023) Influence of pH on indole-dependent heterodimeric interactions between Anopheles gambiae odorant-binding proteins OBP1 and OBP4. Int J Biol Macromol 245: 125422

Mann HB, Whitney DR (1947) On a Test of Whether One of 2 Random Variables Is Stochastically Larger Than the Other. Ann Math Stat 18: 50–60

Manoharan M, Fuchs PF, Sowdhamini R, Offmann B (2014) Insights on pH-dependent conformational changes of mosquito odorant binding proteins by molecular dynamics simulations. Journal of biomolecular structure & dynamics 32: 1742–51

Manoharan M, Ng Fuk Chong M, Vaïtinadapoulé A, Frumence E, Sowdhamini R, Offmann B (2013) Comparative Genomics of Odorant Binding Proteins in Anopheles gambiae, Aedes aegypti, and Culex quinquefasciatus. Genome Biology and Evolution 5: 163–180

Meylan WM, Howard PH, Boethling RS (1996) Improved method for estimating water solubility from octanol water partition coefficient. Environ Toxicol Chem 15: 100–106

Murphy EJ, Booth JC, Davrazou F, Port AM, Jones DN (2013) Interactions of Anopheles gambiae odorant-binding proteins with a human-derived repellent: implications for the mode of action of n,n-diethyl-3-methylbenzamide (DEET). J Biol Chem 288: 4475–85

Murshudov GN, Vagin AA, Dodson EJ (1997) Refinement of macromolecular structures by the maximum-likelihood method. Acta Crystallographica Section D-Biological Crystallography 53: 240–255

Musah RA, Jensen GM, Rosenfeld RJ, McRee DE, Goodin DB, Bunte SWJJotACS (1997) Variation in strength of an unconventional C− H to O hydrogen bond in an engineered protein cavity. 119: 9083–9084

Pan Y, Huang N, Cho S, MacKerell AD, Jr. (2003) Consideration of molecular weight during compound selection in virtual target-based database screening. J Chem Inf Comput Sci 43: 267–72

Pelletier J, Guidolin A, Syed Z, Cornel AJ, Leal WS (2010) Knockdown of a mosquito odorant-binding protein involved in the sensitive detection of oviposition attractants. J Chem Ecol 36: 245–8

Pelosi P, Iovinella I, Zhu J, Wang G, Dani FR (2018) Beyond chemoreception: diverse tasks of soluble olfactory proteins in insects. Biological Reviews 93: 184–200

Pu J, Chung H (2024) New and emerging mechanisms of insecticide resistance. Current Opinion in Insect Science 63: 101184

Qiao H, He X, Schymura D, Ban L, Field L, Dani FR, Michelucci E, Caputo B, Torre AD, Iatrou K, Zhou JJ, Krieger J, Pelosi P (2010) Cooperative interactions between odorant-binding proteins of Anopheles gambiae. Cell Mol Life Sci 68: 1799–1813

Rana A, Sharma D, Choudhary K, Kumari P, Ruchika K, Yangchan J, Kumar S (2024) Insight into insect odorant binding proteins: An alternative approach for pest management. Journal of Natural Pesticide Research 8: 100069

Roberts E, Eargle J, Wright D, Luthey-Schulten Z (2006) MultiSeq: unifying sequence and structure data for evolutionary analysis. Bmc Bioinformatics 7

Sandler BH, Nikonova L, Leal WS, Clardy J (2000) Sexual attraction in the silkworm moth: structure of the pheromone-binding-protein-bombykol complex. Chem Biol 7: 143–51

Sanner MF (1999) Python: a programming language for software integration and development. J Mol Graph Model 17: 57–61

Sarkhel S, Desiraju GR (2004) N H…O, O H…O, and C H…O hydrogen bonds in protein–ligand complexes: Strong and weak interactions in molecular recognition. 54: 247–259

Spinelli S, Lagarde A, Iovinella I, Legrand P, Tegoni M, Pelosi P, Cambillau C (2012) Crystal structure of Apis mellifera OBP14, a C-minus odorant-binding protein, and its complexes with odorant molecules. Insect biochemistry and molecular biology 42: 41–50

Stierand K, Maass PC, Rarey M (2006) Molecular complexes at a glance: automated generation of two-dimensional complex diagrams. Bioinformatics 22: 1710–1716

Sun JS, Xiao S, Carlson JR (2018) The diverse small proteins called odorant-binding proteins. Open biology 8: 180208

Tegoni M, Campanacci V, Cambillau C (2004) Structural aspects of sexual attraction and chemical communication in insects. Trends in Biochemical Sciences 29: 257–264

Thireou T, Kythreoti G, Tsitsanou KE, Koussis K, Drakou CE, Kinnersley J, Krober T, Guerin PM, Zhou JJ, Iatrou K, Eliopoulos E, Zographos SE (2018) Identification of novel bioinspired synthetic mosquito repellents by combined ligand-based screening and OBP-structure-based molecular docking. Insect Biochem Mol Biol 98: 48–61

Trott O, Olson AJ (2010) Software News and Update AutoDock Vina: Improving the Speed and Accuracy of Docking with a New Scoring Function, Efficient Optimization, and Multithreading. Journal of Computational Chemistry 31: 455–461

Tsitsanou KE, Thireou T, Drakou CE, Koussis K, Keramioti MV, Leonidas DD, Eliopoulos E, Iatrou K, Zographos SE (2012) Anopheles gambiae odorant binding protein crystal complex with the synthetic repellent DEET: implications for structure-based design of novel mosquito repellents. Cellular and Molecular Life Sciences 69: 283–297

Tsouri A, Douris V (2025) The Role of Chemosensory Proteins in Insecticide Resistance: A Review. Insects 16

US-EPA (2026) US EPA. [2026]. Estimation Programs Interface Suite™ for Microsoft® Windows, v 4.11. United States Environmental Protection Agency, Washington, DC, USA. In

Vagin A, Teplyakov A (1997) MOLREP: an automated program for molecular replacement. Journal of Applied Crystallography 30: 1022–1025

Venthur H, Zhou JJ (2018) Odorant Receptors and Odorant-Binding Proteins as Insect Pest Control Targets: A Comparative Analysis. Frontiers in physiology 9: 1163

Vogt RG, Riddiford LM (1981) Pheromone Binding and Inactivation by Moth Antennae. Nature 293: 161–163

Wang Q, Shentu X, Yu X, Liu Y (2025) Insect Odorant-Binding Proteins (OBPs) and Chemosensory Proteins (CSPs): Mechanisms and Research Perspectives in Mediating Insecticide Resistance. Biology (Basel) 14

Wishart DS, Feunang YD, Guo AC, Lo EJ, Marcu A, Grant JR, Sajed T, Johnson D, Li C, Sayeeda Z, Assempour N, Iynkkaran I, Liu Y, Maciejewski A, Gale N, Wilson A, Chin L, Cummings R, Le D, Pon A et al. (2018) DrugBank 5.0: a major update to the DrugBank database for 2018. Nucleic Acids Res 46: D1074–D1082

Yin J, Choo YM, Duan H, Leal WS (2015) Selectivity of odorant-binding proteins from the southern house mosquito tested against physiologically relevant ligands. Front Physiol 6: 56

Zheng J, Li J, Han L, Wang Y, Wu W, Qi X, Tao Y, Zhang L, Zhang Z, Chen Z (2015) Crystal structure of the Locusta migratoria odorant binding protein. Biochemical and Biophysical Research Communications 456: 737–742

Zheng J, Yang M, Dong K, Zhang J, Wang H, Xie M, Wu W, Zhang Y-J, Chen ZJIJoMS (2022) Structural insights into the ligand-binding and-releasing mechanism of Helicoverpa armigera pheromone-binding protein PBP1. 23: 1190

Zhou J-J (2010) Chapter Ten - Odorant-Binding Proteins in Insects. In Vitamins & Hormones, Litwack G (ed) pp 241–272. Academic Press

Ziemba BP, Murphy EJ, Edlin HT, Jones DN (2013) A novel mechanism of ligand binding and release in the odorant binding protein 20 from the malaria mosquito Anopheles gambiae. Protein science : a publication of the Protein Society 22: 11–21

Zographos SE, Eliopoulos E, Thireou T, Tsitsanou KE (2018) OBP-structure-aided repellent discovery: An emerging tool towards the prevention of mosquito-borne diseases. In Computational design of chemicals for the control of mosquitoes and their diseases, Devillers J (ed) pp 65–105. CRC press book series "QSAR in Environmental and Health Sciences"

