## Supplementary Figures and Tables for "AgamOBP4-targeted drug repurposing identifies novel mosquito repellent scaffolds"

**Full length amino acid sequence of the mature AgamOBP4**

**>tr|Q8T6R7|26-150**

**1** AMTMKQLTNS MDMMRQACAP KFKVEEAELH GLRKSIFPAN PDKELKCYAM CIAQMAGTMT

**61** KKGEISFSKT MAQIEAMLPP EMKTMAKEAL THCKDTQTSY KDPCDKAYFS AKCAADFTPD

**121** TFMFP

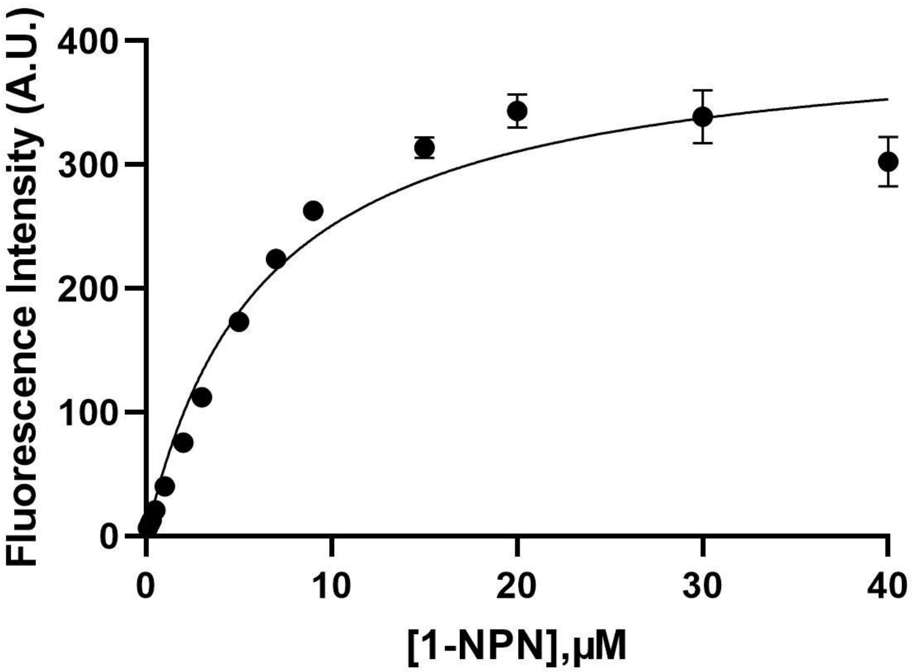


**Figure S1. Saturation binding curve of 1-NPN.** AgamOBP4 (2 µM) was titrated with 1-NPN over a concentration range of 0.1–40 µM. The Kd of 1-NPN for AgamOBP4 was determined to be 6.29 µM. The assay conditions and binding-model equation are described in the Experimental Section.

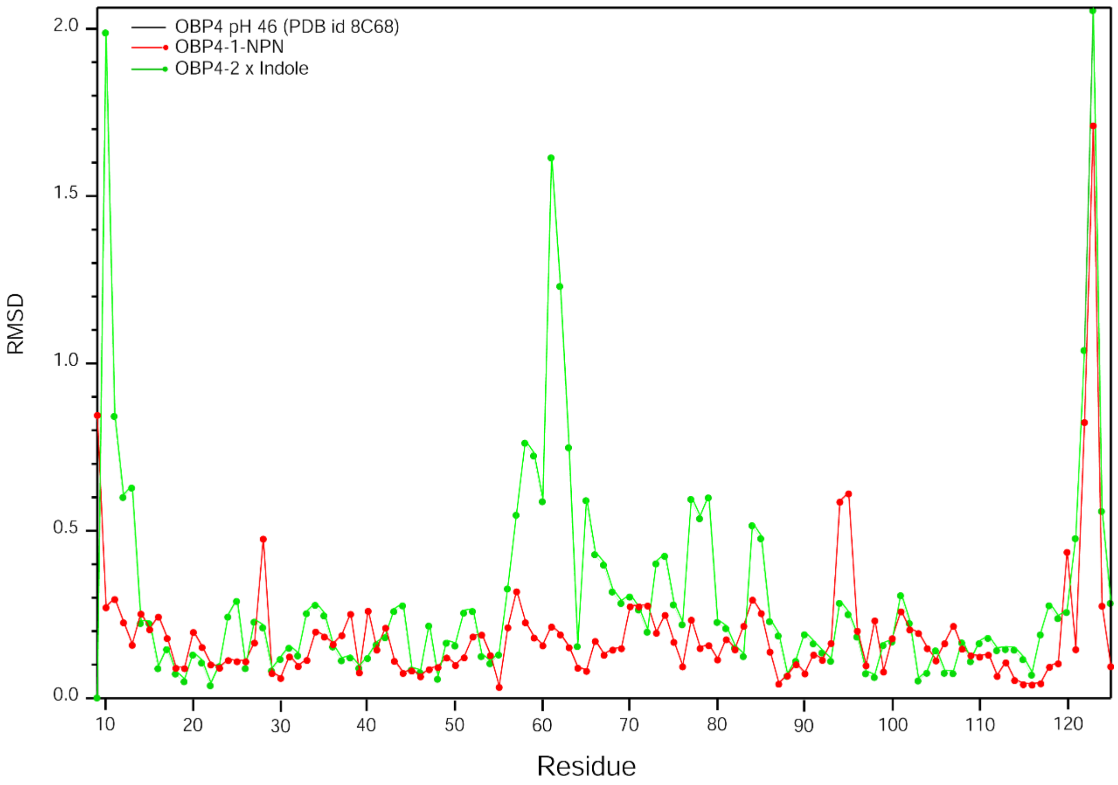


**Figure S2. RMSD vs Residue number**. Plots of the RMSD of the mainchain atom coordinates versus the residue number of AgamOBP4-1-NPN (red), and AgamOBP4-indole at pH 4.6 (green) structures compared to the apo AgamOBP4 at pH 4.6 (PDB ID: 8C68, reference structure).

### Table S1. Protenix confidence metrics for predicted AgamOBP4–ligand complexes

| Ligand | [a]pTM | [b]ipTM | [c]pLDDT | [d]GPDE |
|---|---|---|---|---|
| 1-NPN | 0.95 | 0.86 | 85.88 | 0.47 |
| 2 x Indole | 0.96 | 0.86 | 90.15 | 0.55 |
| 3 | 0.96 | 0.88 | 85.90 | 0.41 |
| 5 | 0.95 | 0.85 | 85.90 | 0.49 |
| 12 | 0.95 | 0.85 | 85.90 | 0.49 |
| 14 | 0.95 | 0.86 | 85.40 | 0.48 |
| 15 | 0.96 | 0.85 | 85.91 | 0.46 |
| 18 | 0.95 | 0.88 | 85.89 | 0.48 |
| 22 | 0.95 | 0.83 | 85.92 | 0.51 |

**[a]pTM (predicted template modelling)** Predicted TM-score for the full predicted complex (0 to 1). Estimates overall topological accuracy of the predicted fold; values above ~0.5 indicate the model likely captured the correct global fold, above ~0.8 high confidence. Typical range: ~0.34 to 0.95.

**[b]ipTM** (**interface predicted template modelling**) Interface predicted TM-score (0 to 1), measuring confidence in the relative positioning of chains at inter-chain interfaces. The primary metric for multi-chain complex / binder confidence; above ~0.6 typically indicates a confidently predicted interface. Typical range: ~0.17 to 0.93.

**[c]pLDDT (The predicted local distance difference test**) Mean predicted Local Distance Difference Test score for the sample, on a 0 to 100 scale (e.g. ~43 to 94, mean ~83). Per-residue confidence in local structure averaged over the model; above 90 very high, 70 to 90 confident, below 50 low confidence / likely disordered.

**[d]GPDE (Global Predicted Distance Error)** for the sample, summarizing the model's expected error in pairwise residue distances. Lower is better. Useful as a complementary confidence signal to pTM/ipTM. Typical range: ~0.36 to 2.76 (not bounded to 0-1).

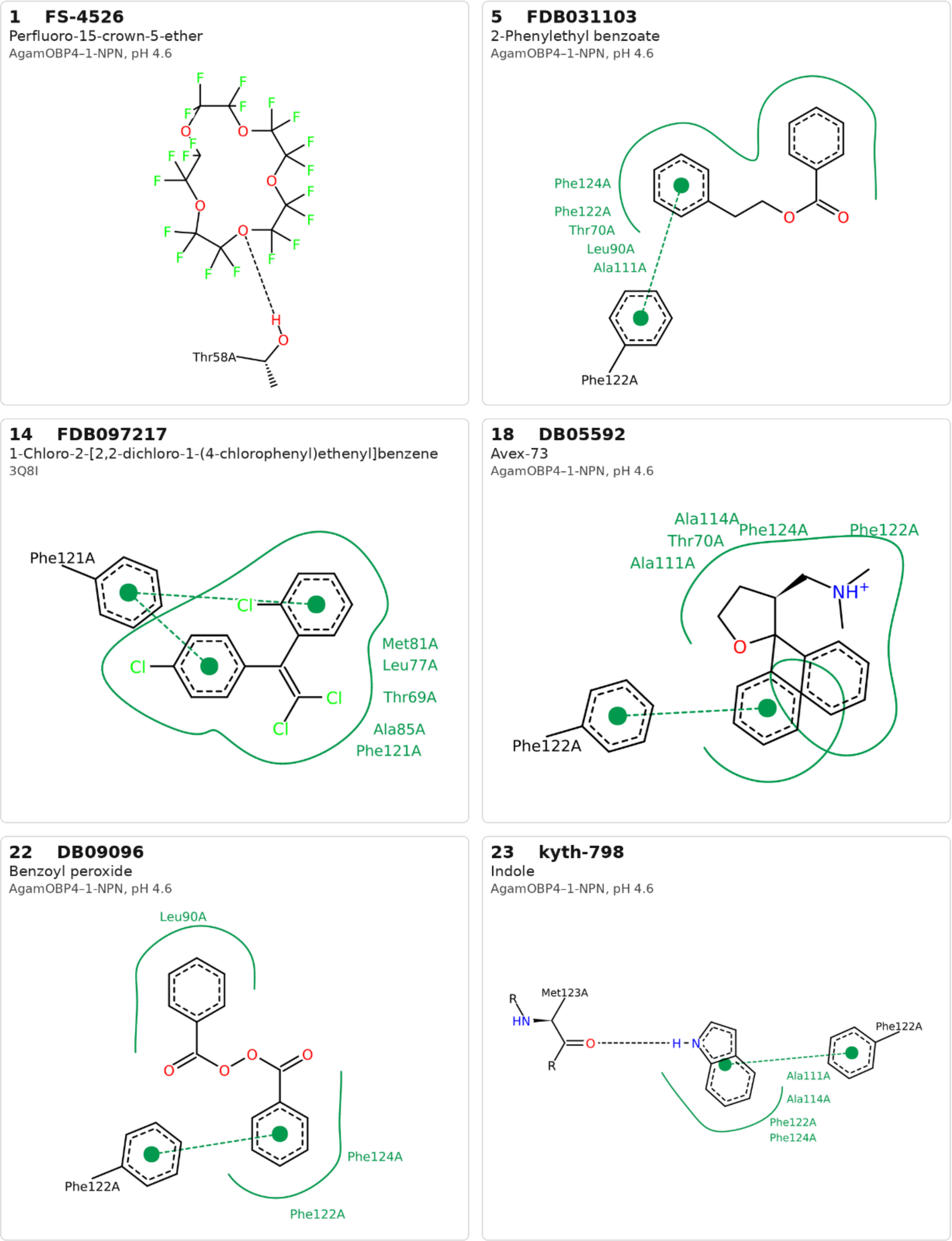


**Figure S3.** Visual representation of the binding poses of selected compounds in AgamOBP4 crystals, generated with PoseView. All shown poses are for the protein in which the compounds had the best pose.

**Table S2.** Protenix confidence metrics for the predicted AgamOBP4-Fluorene complexes

| **Fluorene** | $^{a}$**pTM** | $^{b}$**ipTM** | $^{c}$**pLDDT** | $^{d}$**GPDE** |
|---|---|---|---|---|
| **Pose 0** | 0.959 | 0.879 | 85.897 | 0.411 |
| **Pose 1** | 0.959 | 0.878 | 85.897 | 0.413 |
| **Pose 2** | 0.958 | 0.876 | 85.894 | 0.413 |
| **Pose 3** | 0.959 | 0.875 | 85.901 | 0.413 |
| **Pose 4** | 0.959 | 0.874 | 85.897 | 0.414 |

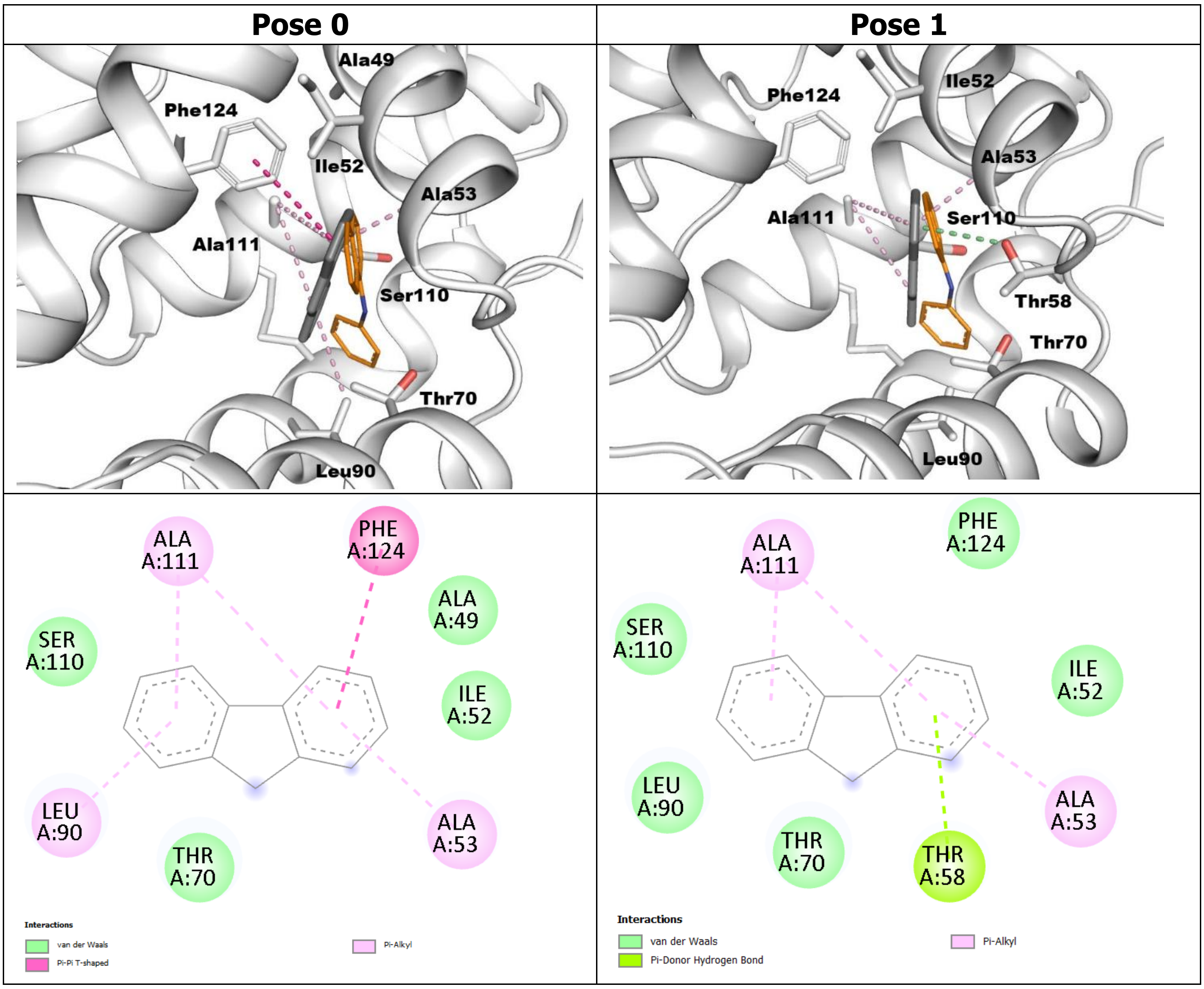


**Figure S4. The two top ranked poses of Fluorene (#3).** The top ranked pose (Pose 0) lacks any of the binding mode criteria proposed for repellent activity. In the second top ranked pose (pose 1), fluorene is predicted to approximate crystallographic molecules and to form a π-donor hydrogen bond between the aromatic ring in indole-1 subsite. Furthermore, the CG2 atom of Thr70 lies in close proximity to C1, C2, and C3 carbon atoms of the fluorene moiety in the indole-2 subsite (3.37, 3.50, and 3.69 Å, respectively).
